# Structural underpinning of multi-functionality of TFIIE-related tandem-winged-helix in RNA polymerase III

**DOI:** 10.1101/2024.08.27.609947

**Authors:** Jheng-Syong Wu, Yu-Chun Lin, Yi-Yu Wei, Hsin-Hung Lin, Yang-Chih Liu, Jen-Wei Chang, I-Ping Tu, Hung-Ta Chen, Wei-Hau Chang

## Abstract

The tandem-winged-helix (tWH) domain of RNA polymerase III (Pol III) Rpc34 subunit is a multi-function hub in Pol III apparatus while the structure basis remains elusive. To probe tWH in Pol III elongation complex (EC), we engineered an azide-bearing unnatural amino acid into tWH and implemented a thiol-capping scheme to enable strain-promoted click reaction for selective dye labeling to suit single-molecule Förster resonance energy transfer (smFRET). Our discrete smFRET results and nano-positioning analysis reveal multiple docking sites of Rpc34 WH2 on Pol III EC with characteristic dwell-times, reflecting promiscuous and weak interactions underpinning tWH’s multi-functionality. The docking sites include one overlapping with that during initiation and others on transcription bubble and downstream DNA, previously unreported. This work provides mechanistic insights into Pol III transcription re-initiation and elongation, with useful bio-orthogonal strategies for studying structural dynamics of large native protein complexes.

## Introduction

Among three eukaryotic nuclear RNA polymerases, RNA polymerase III (Pol III) is a highly efficient enzyme specialized for synthesizing structural and non-coding RNAs that are essential for eukaryotic cells. Pol III-synthesized RNAs are typically short, only a few hundred nucleotides long, and include tRNAs, 5S rRNA, U6 snRNA, and 7SL RNA, among others^1,2^. The quantity of Pol III-synthesized RNAs surpasses the sum of all other types of RNA in the cell. This high output necessitates that Pol III to initiate and terminate transcription more frequently than Pol I and Pol II. In humans, dysregulation of these RNAs due to defects in the Pol III transcription machinery is linked to a variety of diseases, including neurodegenerative disorders and several types of cancer^3–5^.

The Pol III machinery consists of 17 subunits, forming a complex with a molecular weight of approximately 0.7 MD. Presumably evolved from Pol II, Pol III acquired five additional subunits^2^, which are organized into two sub-complexes: the Rpc82/34/31 hetero- trimer and the Rpc53/37 hetero-dimer^2^. These Pol III-specific subunits are related to the Pol II initiation factors TFIIE and TFIIF^6,7^, and thus appear to play roles exclusively in Pol III initiation. However, unlike TFIIE and TFIIF, which transiently associate with Pol II during initiation, these sub-complexes remain stably and permanently associated with Pol III as integral components. This suggests they may also function as “built-in” elongation factors, contributing to elongation, termination, or re-initiation^2^. In fact, they may hold the keys to the high efficiency of Pol III transcription.

Among the Pol III-specific subunits in yeast, Rpc34 (known as RPC6 in humans) is unique component—it is within the TFIIE-related heterotrimeric sub-complex, and it lacks a counterpart in the other two nuclear RNA polymerases, Pol I and Pol II. Rpc34 is essential to Pol III transcription despite it is a small protein composed of 317 amino acids. Early functional studies in yeast have revealed Rpc34’s dual roles in Pol III transcription initiation^8^, in opening promoter DNA and mediating the formation of the initiation complex (IC) ^9^, respectively. The amino acids responsible for these activities are located in the protein’s N-terminal region, which adopts a tandem winged-helix (WH) fold^10^, comprising WH1 (amino acids 12 to 78) and WH2 (amino acids 90 to 151), collectively referred to as the tandem WH (tWH) domain. Remarkably, recent genetic studies have revealed Rpc34’s tWH domain also plays roles in the stage of elongation^11^. These findings, together, indicate that Rpc34 tWH is a multi-function domain and suggest it may engage in different functions during different stages of transcription.

To explore the structural basis of Rpc34 tWH’s function(s), the photo-activated crosslinking (PA-XL) technique^12^ was employed prior to the advent of high-resolution cryo-electron microscopy (cryo-EM). This approach allowed for the provisional placement of Rpc34 tWH above the Pol III DNA-binding cleft in the upstream region (Supplementary Fig. 1), aligning with its role in opening promoter DNA during initiation. Strikingly, subsequent high-resolution cryo-EM structures of the Pol III initiation complex (IC) ^13,14^ revealed that Rpc34 tWH could shift further upstream along the DNA-binding cleft, towards the Rpc160 (Rpc1) protrusion domain (Supplementary Fig. 1). At this new position, Rpc34 WH2 domain literally contacts the protrusion domain and the promoter-bound initiation factor TFIIIB, mediating the formation of IC. Notably, this site can also be alternatively occupied by Maf1, resulting in suppression of Pol III transcription^15^ (Supplementary Fig. 1).

As Pol III transcription transitions to the elongation stage, the structural basis for Rpc34 tWH’s function(s) becomes obscure since the Rpc34 tWH density is missing in the cryo-EM map of Pol III elongation complex (EC)^16^. This absence implies that Rpc34 tWH is highly dynamic, but the definitive evidence is lacking. Importantly, this presumed structural dynamics is likely to be a conserved feature from yeast to human, as the tWH domain of RPC6/RPC39 (the human homolog of Rpc34) is also missing in the corresponding cryo-EM structure^17^.

To investigate the dynamics of Rpc34 tWH in the Pol III elongation complex (EC), we utilized single-molecule Förster resonance energy transfer (smFRET), an imaging-based method sensitive to conformation dynamics at molecular scale and capable of uncovering information masked by ensemble measurements^18^. When applied to immobilized molecules, smFRET tracks time-trajectories of individual molecules, allowing for extraction of kinetics information without requiring synchronization. However, using smFRET to study large protein complexes faces a significant challenge at the level of site-specific fluorophore labeling. Standard site-specific protein labeling typically employs nucleophile-reactive maleimides to target cysteine residues, which requires editing naturally occurring cysteines. This approach is unsuitable for Pol III, as it contains more than 60 naturally occurring cysteine residues, constituting approximately 1% of its total amino acids (6,600). To overcome this limitation, we adopted a bio-orthogonal approach using unnatural amino acids (UAAs)^19,20^. We modified our previously developed yeast genetic system used for incorporating a photoactive UAA into a designated site in RNA polymerases^12,21^ for introducing a different UAA suitable for site-specific fluorophore labelling using click chemistry. While this UAA-based site-specific labelling strategy has been demonstrated in a number of smFRET studies, its application has so far been limited to small test proteins^22,23^, or simple protein complexes assembled in vitro^24^. Thus, whether this approach that hinges on “clicking” a UAA can be readily applied to large or native protein complexes such as Pol III remains to be determined.

In this study, we site-specifically incorporated azido-phenylalanine (Fig. 1) into Rpc34 WH2 and conjugated it with an alkyne-bearing fluorophore (DIBO-dye) using strain-promoted azide-alkyne [3+2] cycloaddition (SpAAC), best known as Bertozzi’s click reaction^25^. This straightforward click chemistry is biocompatible as it avoids the use of hazardous copper^25^, making it suitable for various bio-orthogonal applications. To our dismay, our bulk labeling experiments revealed extensive labeling of numerous Pol III subunits that had not been modified with the UAA (azido-phenylalanine). To address this, we developed a simple chemical step to suppress cross-reactivity between cyclooctynes and thiols^26,27^, successfully enabling selective dye labeling of the UAA (azido-phenylalanine) incorporated into Pol III.

**Fig. 1.**
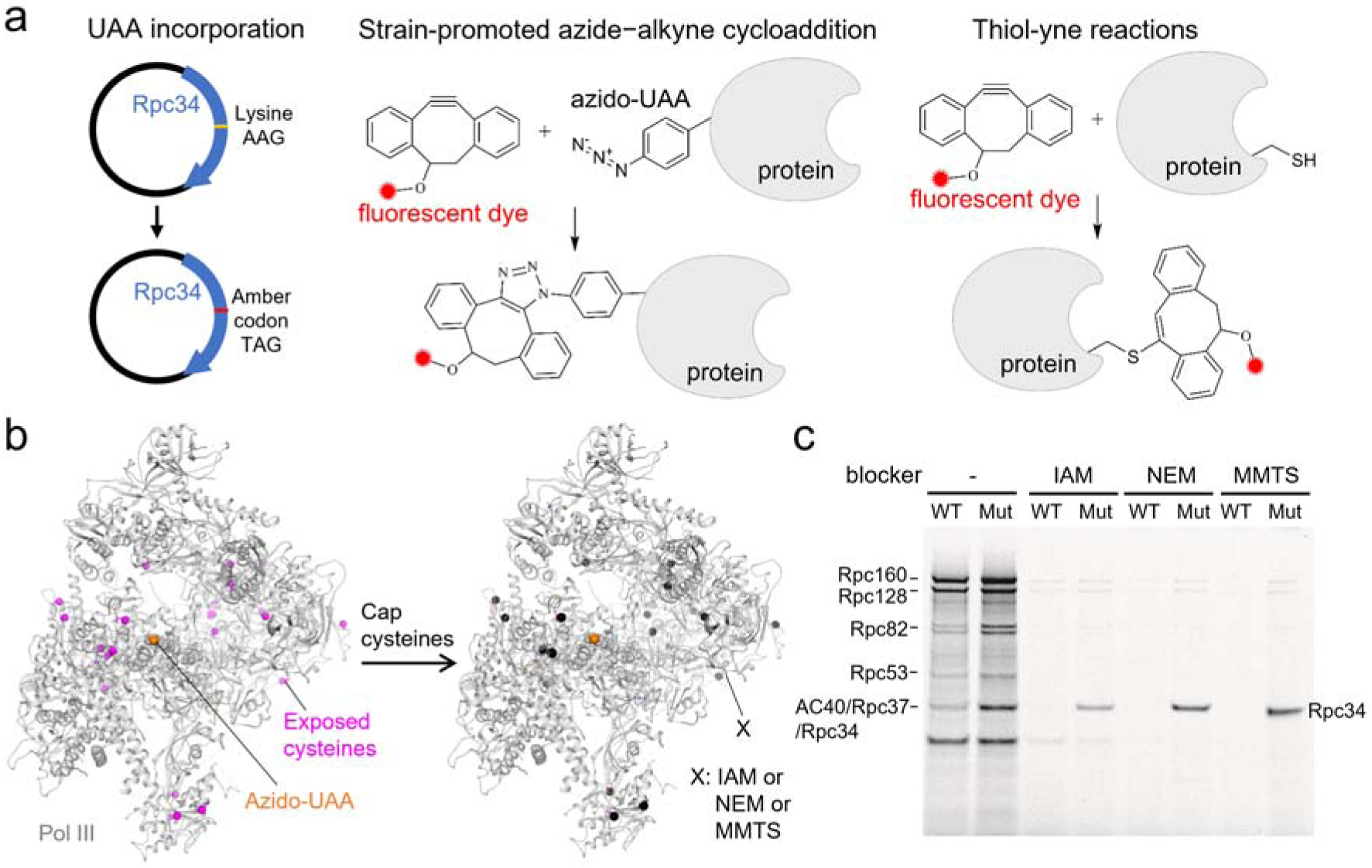
Selective Dye Labeling of Incorporated Azido-UAA by Capping Native Cysteines in Pol III. **(a)** *Left:* Schematic representation of unnatural amino acid (UAA) incorporation into Rpc34 WH2 in Pol III. The Rpc34/pRS425 plasmid was transformed into a yeast strain lacking chromosomal Rpc34, where a specific codon was mutated to an amber codon for the incorporation of azido-UAA at a designated position. *Middle:* Chemical structures of DIBO (dibenzocyclooctyne, a strained alkyne) and azido-phenylalanine UAA on a protein ready for labeling via strain-promoted azide-alkyne [3+2] cycloaddition (SpAAC). *Right:* The reaction of thiol-yne addition, where free thiol groups on cysteines react with the alkyne-dye, leading to non-specific DIBO dye labeling on Pol III. **(b)** Cysteine Blocking Schematic: This illustration shows that exposed cysteines (magenta spheres) on Pol III can be capped by thiol-reactive molecules (black spheres), inhibiting non-specific DIBO dye labeling. **(c)** Screening of Thiol Blockers: Thiol-reactive molecules—iodoacetamide (IAM), N- ethylmaleimide (NEM), and S-methyl methanethiosulfonate (MMTS)—were tested at a concentration of 20 mM. The Alexa647 fluorescence image of an SDS-PAGE gel shows that IAM, NEM, and MMTS effectively abolished non-specific DIBO-Alexa647 labeling of wild- type Pol III and azido- UAA-modified Pol III (see also Supplementary Fig. 2).

Having labeled the azido-UAA incorporated in Rpc34 WH2 with the acceptor dye (an alkyne dye), we measured smFRET between the acceptor and the donor on a site in DNA in a Pol III EC assembled from Pol III, and DNA/RNA scaffold (Fig. 1). Our smFRET results revealed multiple discrete FRET states and captured inter-state transitions, compared with a single FRET state observed when Rpc34 WH2 was constrained by Maf1. Further nano- positioning analysis^28,29^ based on FRET distance constraints localized multiple docking sites of Rpc34 WH2 on EC with spatial precision better than 10 Å. Those findings uncover dynamic positioning of a domain in the TFIIE-related sub-complex in Pol III, underpinning Rpc34 tWH multi-functionality, with the first glimpse of Pol III as dynamic apparatus to provide molecular insights into the highly efficient Pol III transcription.

## Results

### Capping cysteines ensures selective SpAAC-based UAA-dye conjugation

To incorporate 4-Azido-L-phenylalanine (AzF) into Rpc34 within native Pol III, we modified our yeast UAA system previously developed for the incorporation of benzoyl-phenylalanine (BPA) ^12,21^ (Fig. 1a & **Methods**). We then site-specifically substituted a Rpc34 residue with AzF, and tested various single amino acid mutants substituted with AzF — none of them exhibited a detectable phenotype. Successful incorporation of AzF was confirmed by the cell growth dependency on AzF following the genetic modification of Rpc34. We purified both wild-type Pol III and the Rpc34 mutant bearing AzF, and labeled those enzymes with DIBO-Alexa-647 (DIBO, dibenzocyclooctyne, a strained alkyne) using the click chemistry of strain-promoted alkyne cyclooctyne addition (SpAAC), commonly referred to as copper-free “Bertozzi” click reaction^25^ (Fig. 1a). So far, only a limited number of smFRET studies have utilized this labeling scheme, and those studies have been restricted to small test proteins^22,23^ or simple protein complexes assembled in vitro from recombinant proteins^24^. Thus, the feasibility of the Bertozzi labelling approach for large endogenous protein complexes like Pol III remains to be determined. Interestingly, while the DIBO-dye labeling was compatible with Pol III transcription activity (data not shown), it yielded extensive dye labeling of Pol III subunits that had not been substituted with AzF (see *lane* 1, Fig. 1c). These “non-specifically” labelled subunits included Rpc1 (Rpc160), Rpc2 (Rpc128), Rpc82, Rpc37/AC40, and ABC27. Given that these subunits possess abundant native cysteines (Supplementary Table 1), we hypothesized that the unwanted labeling likely stemmed from surface-exposed cysteines reacting with the alkyne moiety in DIBO (Fig. 1a). This hypothesis is supported by a previous report indicating that azide-dependent protein labeling via strain-promoted alkyne-azide cyclooctyne (SpAAC) addition was compromised by thiol-ene addition reactions^26,27^.

To address this issue, we explored the use of small molecules that could cap the thiol groups^30^ of exposed cysteines in Pol III (Fig. 1b). We tested several compounds— iodoacetamide (IAM), N-ethylmaleimide (NEM), and methyl-methanethiosulfonate (MMTS)—and evaluated their impact on labeling AzF-substituted Pol III compared to wild-type Pol III as a control. As shown in Fig. 1c (*lanes* 3-8), all tested thiol-capping molecules effectively abolished unwanted labeling, which in turn achieved selective labeling of the AzF-substituted Rpc34 subunit in Pol III. Since IAM was found to also significantly reduce specific Rpc34 labeling compared to NEM or MMTS, we dispensed its further use. We further optimized the concentrations of NEM or MMTS (Supplementary Fig. 2) to avoid interference with Pol III EC formation.

### SmFRET reveals Rpc34 WH2 is highly mobile

To probe the dynamics of Rpc34 WH2 in the Pol III elongation complex (EC) using single-molecule Förster resonance energy transfer (smFRET), we first selected the K126 residue in Rpc34 for acceptor dye labeling, with the donor dye placed at DNA(+7) (Fig. 2a), a site located slightly downstream of the active center and exhibiting high structural order in the cryo-EM structure^16^. Before investigating Rpc34 WH2 dynamics in Pol III-EC, we validated the precision of our smFRET system through a benchmark DNA test^35^ (Supplementary Fig. 3), and aimed to establish a control experiment for static smFRET when the mobility of Rpc34 WH2 is restrained. Inspired by Rpc34 WH2 stabilized by Maf1 in the cryo-EM structure of the Pol III-Maf1 complex^15^, we used Maf1 to restrict the mobility of Rpc34 WH2 in the control experiment. As Rpc34 WH2 is situated at this position^15^ (Fig. 2b), the distance between the centers of the simulated swirling dye volume^31^ of the acceptor dye labeled on K126 and that of the donor dye attached to DNA(+7) is 7 nm^31^. This distance yielded a low FRET efficiency (E_model_) of 0.26 for the dye pair of TAMRA and Alexa647, which has a Förster distance of 6 nm^32,33^.

**Fig. 2.**
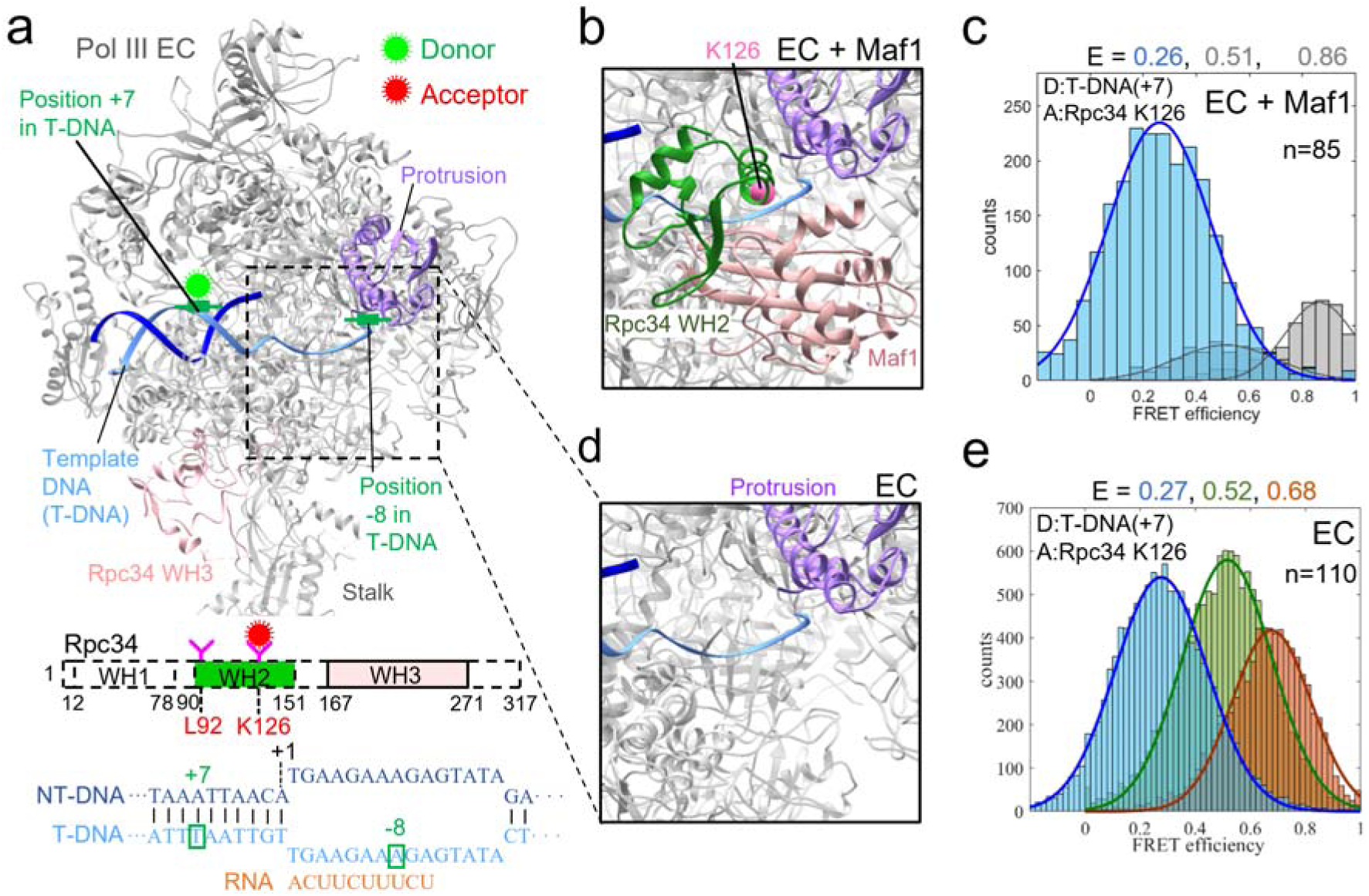
Pol III EC Structure and smFRET Experiments. (**a**) The design of the donor position for the FRET pair is based on the yeast Pol III elongation complex (EC) structure (PDB: 5fj8, gray), where the cryo-EM density of the Rpc34 tWH domain is missing. For clarity, the Rpc82 and Rpc31 subunits have been computationally removed from the displayed structure. The donor dye (TAMRA) is labeled at the +7 position of the template DNA (T-DNA, cyan), while the acceptor dye (Alexa647) is attached to an amino acid in Rpc34 WH2, such as K126. (**b**) Rpc34 WH2 is stabilized by the Maf1 protein (PDB: 6tut). (**c**) The smFRET histogram from a Pol III EC experiment with Maf1 added shows a dominant population centered at a mean FRET efficiency of 0.26 (E: mean FRET efficiency, n: number of molecules). (**d**) The Rpc34 WH2 domain is absent in the Pol III-EC structure (PDB: 5fj8, gray). (**e**) The smFRET histogram from a Pol III EC experiment conducted in the absence of Maf1 is resolved into three distinct populations: low FRET (light blue), middle FRET (light green), and high FRET (orange). Each population is fitted with a single Gaussian function, revealing mean low-, middle-, and high-FRET efficiencies of 0.27, 0.52, and 0.68, respectively.

We then added an overwhelming excess (10× more) of Maf1 to donor-and-acceptor-labeled Pol III EC molecules immobilized on cover slips and performed smFRET experiments using our custom-built total internal reflection fluorescence microscope equipped with alternating laser excitation (ALEX) capability (see **Methods**). In the control experiment with Maf1, we collected a total of 85 time traces exhibiting clear acceptor photo- bleaching signatures^34^ that allowed for extracting absolute FRET efficiencies. As anticipated, nearly all the molecules displayed stationary time trajectories contaminated by shot noise, yielding a mean FRET efficiency (E_measure_) of 0.26 as shown in the frame-wise histogram (Fig. 2c). The excellent agreement between E_measure_ and E_model_ indicates the reliability of our smFRET system as it can recapitulate structural results from cryo-EM^15^. We noticed there were a few molecules with FRET values greater than 0.5, indicating that in some Pol III-EC complexes Maf1 molecules had dissociated from the complexes during the experiments. In addition, we next repeated the control experiment with the acceptor dye attached to L92—

the majority of the molecules exhibited stationary FRET, where the mean FRET efficiency was found to be 0.53 (Supplementary Fig. 4b), in agreement with that from simulation^31^.

Next, we conducted smFRET experiments on donor-acceptor-labeled Pol III EC molecules without Maf1. In the absence of Maf1, smFRET no longer exhibited a single state but revealed various states. In one experiment, we collected a total of 110 time traces; nearly 50% of the molecules (48) exhibited stationary FRET levels that could be roughly classified into high, middle, or low categories, as illustrated by three representative FRET time traces (Supplementary Fig. 4d), with mean efficiencies of 0.7, 0.5, and 0.25, obtained using a wavelet de-noising approach^36^. Notably, the remaining 62 molecules displayed active switching among these FRET levels. The frame-wise histogram (Fig. 2e), constructed from these 110 time traces, clearly resolved the FRET states into three distinct populations, each well fit with a Gaussian distribution.

This analysis allowed for extraction of three mean FRET efficiencies: high (E=0.68), middle (E=0.52), and low (E=0.27). These discrete FRET states with significantly different mean FRET efficiencies indicate that the proximity of Rpc34 WH2 to a fix point in Pol III EC, e.g., DNA(+7), varies vastly. We termed Rpc34 WH2 docking positions corresponding to E=0.27, 0.52 and 0.68 as distal, middle, and proximal sites as reviewed from DNA(+7), where their precise locations warrant further investigation. It is noted that similar dynamic behaviour of Rpc34 was observed when the acceptor was labelled at L92 (Supplementary Fig. 4 & 5), affirming that the observed dynamics of Rpc34 WH2 is regardless of the labelling position.

### Dwell-time analysis reveals characteristic lifetime of each FRET-state

The smFRET time traces exhibiting inter-state switching reflect real-time movement of the Rpc34 WH2 domain. To further characterize the Pol III EC molecules actively switching configurations, we conducted a dwell-time analysis for these FRET states. We employed hidden Markov modeling (HMM) analysis^37^ to identify the time points at which a FRET level change occurs along a time trace, allowing for determining the duration of each FRET level between changes (see **Methods**). With the assistance of alternating laser excitation (ALEX) imaging (Supplementary Fig. 6), we could detect and exclude any false FRET periods caused by acceptor quenching. The pruned time traces were subjected to HMM analysis, revealing high, middle, and low FRET states (Fig. 3a), with estimated FRET efficiencies consistent with those extracted from the frame-wise histogram analysis. The HMM-extracted time durations for each individual FRET state were pooled to create a histogram plotting event numbers against time durations. Remarkably, the distribution in each histogram was well described by a mono-exponential function, yielding defined lifetimes for each state (Fig. 3b). We verified the reproducibility of this observation through three biological repeats (Supplementary Fig. 7), which provided consistent quantitative results: the lifetimes for the high, middle, and low FRET states were determined to be 9.37 ± 1.18, 3.12 ± 0.48, and 1.41 ± 0.11 seconds, respectively, with the high FRET state exhibiting the longest lifetime.

**Fig. 3.**
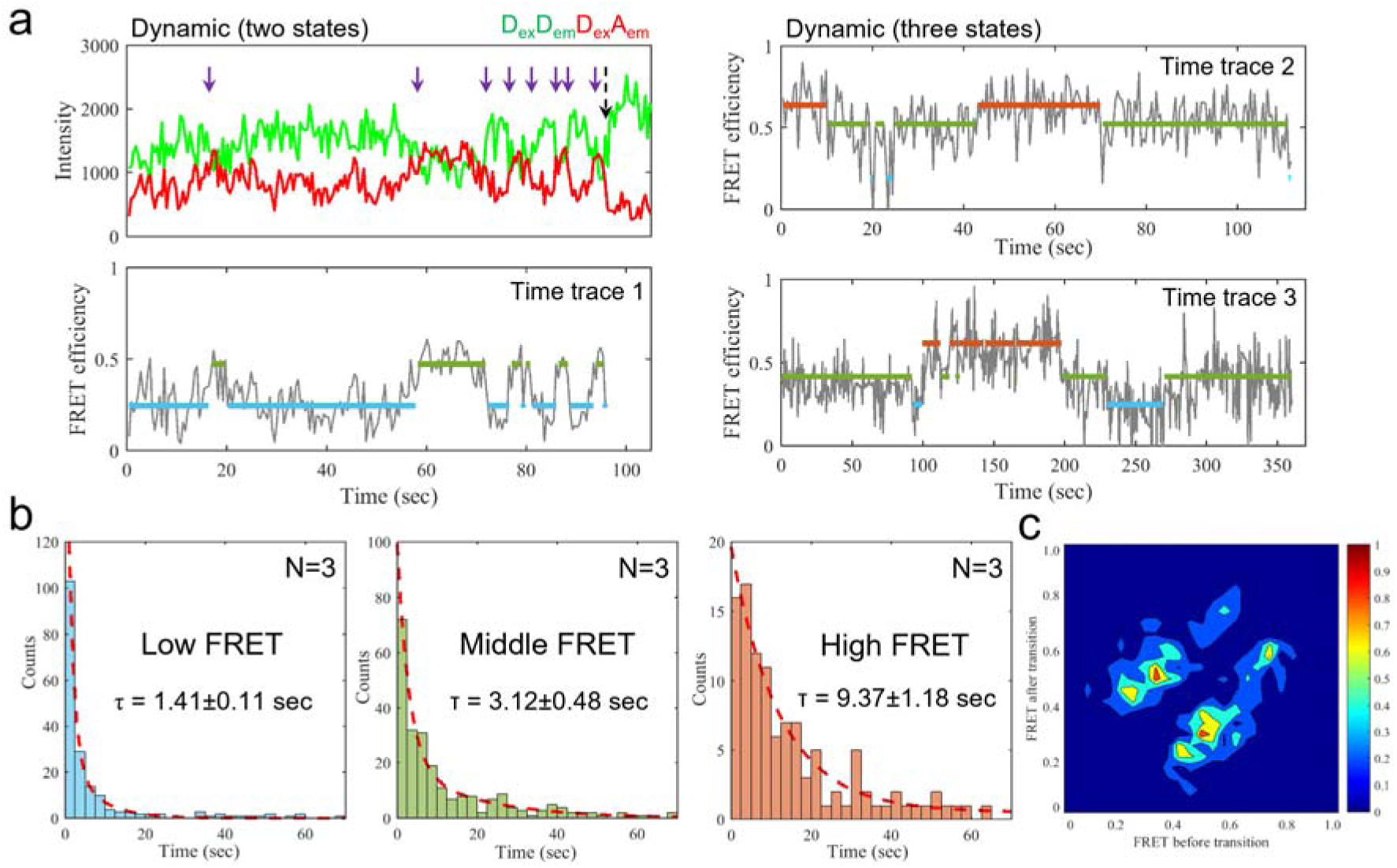
Representative Time Trajectories of Inter-state Converting FRET and Dwell-Time Analysis for Rpc34 WH2 in Pol III EC. The donor dye (TAMRA) and acceptor dye (Alexa647) were labeled at the +7 position of the template DNA and Rpc34 K126, respectively. (**a**) *Left*: A representative time trajectory exhibiting inter-state conversion between two states is shown. The green and red traces represent the donor emission (D_em_) and acceptor emission (A_em_) intensities upon donor excitation, respectively. Purple-black arrows indicate FRET-associated anti-correlated signals between the donor and acceptor, with the last arrow in black highlighting the anti-correlation event resulting from acceptor photo-bleaching. The corresponding FRET time traces (gray) were calculated from the donor and acceptor intensity time traces based on individual γ normalization. Hidden Markov modeling (HMM) was applied to these intensity time traces to identify the levels of each FRET state: low (light blue), middle (light green), and high (orange), along with their respective durations. *Right*: Another two representative time trajectories demonstrate inter-state conversion among three states, with HMM-extracted FRET levels color-coded accordingly (low: light blue, middle: light green, high: orange). (**b**) The dwell-time histograms for low-, middle-, and high-FRET states are presented, based on HMM results from 62 state-changed traces in one experiment. The average dwell time and standard deviation were calculated from three separate experiments (see Supplementary Fig. 7). (**c**) A transition density plot constructed from 633 transition events pooled from multiple experiments illustrates the major and minor populations of transitions. The major transitions occur between the middle-FRET state and the low-FRET state, while minor transitions occur between the middle-FRET state and the high-FRET state.

To explore whether there are preferred transitions among these states, we performed a two-dimensional histogram analysis (Fig. 3c). This analysis indicated that direct transitions between the low and high FRET states appear to be forbidden; instead, every transition involves the middle FRET state. By separating the middle state dwell time into the middle-to-high and the middle-to-low dwell times (Supplementary Fig. 8), we obtained the rate constant for each transition.

### Nano-positioning with smFRET data identifies Rpc34 WH2 docking sites

As mentioned earlier, the high, middle and low FRET states observed for the acceptor labelled at Rpc34 WH2 K126 correspond the proximal, middle and distal site in reference to DNA(+7). To determine spatial locations of those sites within the Pol III EC structure, we employed a nano-positioning system (NPS) analysis^28,29^, a low-resolution structural determination method that can position an antenna in space based on known sites (satellites) using triangulation with FRET distance constraints. This method has been previously used in conjunction with smFRET to elucidate mobile structural elements in protein complex such as RNA polymerase II initiation complex^33^.

Our design of the satellite-antenna (S/A) pairs is illustrated in Fig. 4a. We utilized two satellites: satellite_1 at DNA(+7) in the downstream DNA, and satellite_2 at DNA(-8) in the upstream DNA, to target an antenna of dye labeled at K126 or L92 in Rpc34 WH2. Since L92 is positioned opposite to K126 in the Rpc34 WH2 structure, Rpc34 WH2 can be localized in the region encompassed by the two antennas. The multiple S/A pairs being utilized for this purpose are shown in Fig. 4a. For simplicity, we introduce the notation of S/A to describe a satellite-antenna pair. As an example, +7/126 represents the satellite of the donor dye at DNA (+7) and the antenna of the acceptor dye at Rpc34 K126, while +7/126 =0.26 means this pair gives a mean FRET efficiency of 0.26. To establish a reliable NPS system, we again considered the scenario where Rpc34 WH2 is restrained by Maf1. From a control experiment when Rpc34 WH2 was restrained at the distal site by Maf1, we obtained (+7/126=0.26, -8/126=0.55) (see Fig. 2c for +7/126, and Supplementary Fig. 4c for -8/126). We thereby obtained the NPS credible volume for the antenna dye attached to Rpc34 K126 (Fig. 4c) for positioning Rpc34 WH2 at the distal site. Of note, this experimentally determined NPS credible volume is in good agreement with that obtained from simulation^31^, with a minimal mismatch of 6.9 Å (Supplementary Fig. 9).

**Fig. 4.**
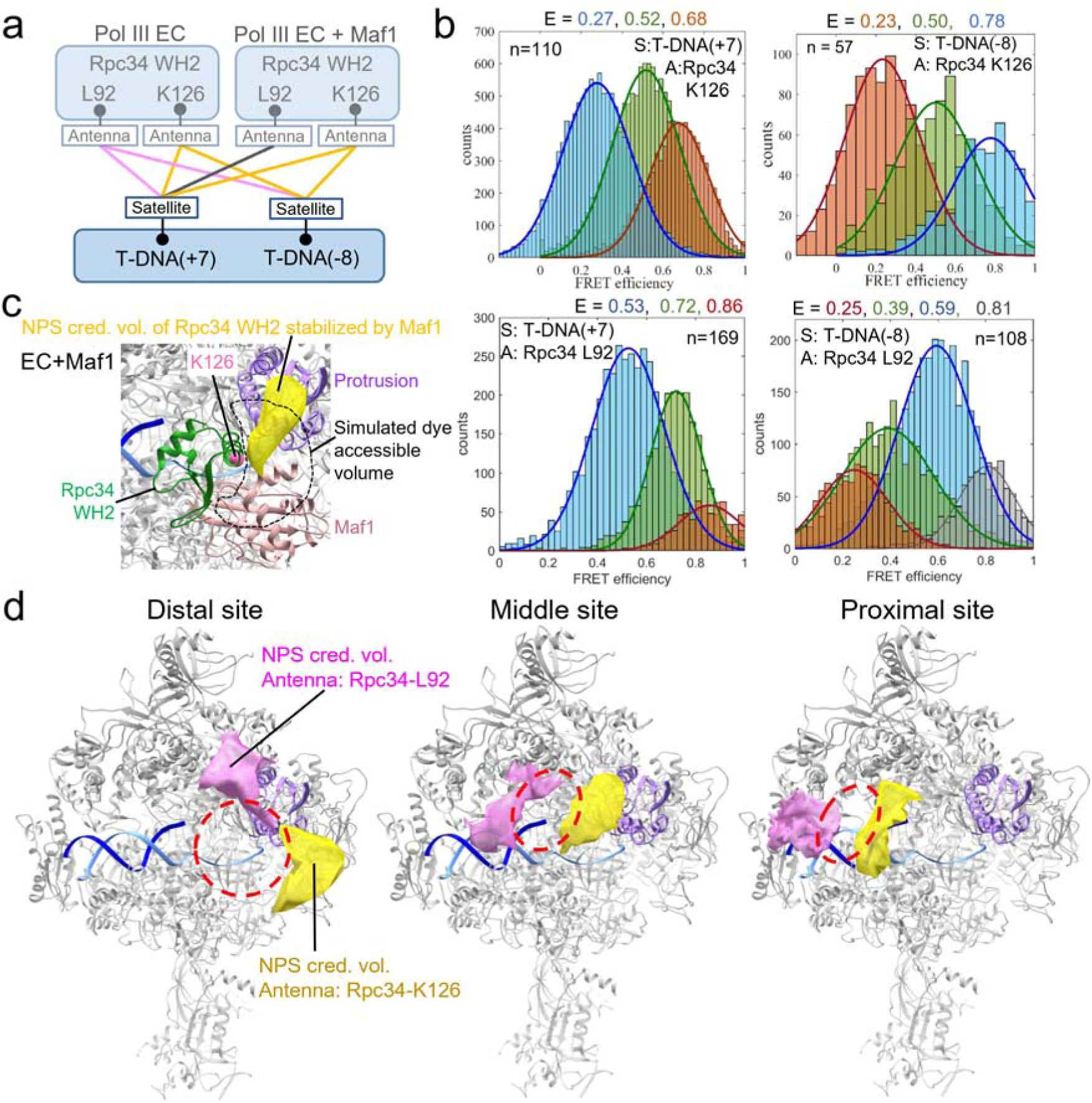
**Nano-Positioning (NPS) Localization of Low-, Middle-, and High-FRET States of Rpc34 WH2 in Pol III EC**. (**a**) Schematic Representation of the FRET Network: This panel illustrates the FRET network utilized for localizing Rpc34 WH2 within the Pol III EC. (**b**) FRET Efficiency Histograms: The histograms depict FRET efficiencies for four satellite-antenna (S/A) pairs: (+7/126, - 8/126) and (+7/92, -8/92), aimed at localizing K126 and L92 of Rpc34. The notation +7/126 indicates that the satellite dye (TAMRA) is labeled at the +7 position in the template DNA (T-DNA), while the antenna dye (Alexa647) is labeled at Rpc34 K126. The colors in the histograms—light blue, light green, and orange—correspond to the low-, middle-, and high-FRET states from the S/A pair of +7/126. The labels E, S, A, and n represent the center of the FRET efficiency distribution, satellite, antenna, and the number of molecules, respectively. (**c**) NPS Localization with Maf1 Stabilization: This panel displays the NPS localization of Maf1-stabilized Rpc34 WH2 in Pol III EC, utilizing two S/A pairs +7/126 and -8/126 (Fig. 2c & Supplementary Fig. 4b). The NPS credible volume (yellow) that localizes the antenna attached to K126 in Maf1-restrained Rpc34 WH2 closely resembles the simulated dye volume (Supplementary Fig. 9). (**d**) NPS Localization of Rpc34 WH2 in Pol III EC: Here, we present the NPS localization of Rpc34 WH2 within Pol III EC with red dashed circles flanked by two credible volumes: one from Rpc34 L92, obtained from the S/A pairs (+7/92, -8/92), and the other from K126, obtained from (+7/126, -8/126).

To localize Rpc34 WH2 without Maf1 stabilization, we first placed the antenna of acceptor dye at its K126 residue, and obtained smFRET results for +7/126 and -8/126 shown in the upper panel of Fig. 4b, from which three meaningful NPS credible volumes (yellow in Fig. 4d) based on the combinations (+7/126=0.27, -8/126=0.78), (+7/126=0.52, -8/126=0.50), and (+7/126=0.68, -8/126=0.23) (Supplementary Fig. 4e), respectively. We next placed the antenna of acceptor dye at L92 to obtained the smFRET results (lower panel Fig. 4b), also yielding three meaningful NPS credible volumes (pink in Fig. 4d) based on the combinations (+7/92=0.53, -8/92=0.59), (+7/92=0.72, -8/92=0.39), and (+7/92=0.86, -8/92=0.25) (Supplementary Fig. 4e), respectively. By identifying the regions between the two dyes attached to Rpc34 L92 and K126, we successfully localized three docking sites of Rpc34 WH2 within Pol III EC (Fig. 4d)—the distal site is situated next to the protrusion domain, in good overlap with the position during transcription initiation^13,14^; whereas the proximal and middle sites are located further downstream along the DNA-binding channel.

## Discussion

In this study, we leveraged single-molecule Förster resonance energy transfer (smFRET) with nano-positioning analysis, a non-canonical structural biology approach capable of tackling dynamic components in a molecular machine^32,33^, to reveal dynamic positioning of the “invisible” Rpc34 tWH domain in the TFIIE-related sub-complex of a Pol III elongation complex. Previous cryo-EM efforts to address the TFIIE-related sub-complex of a Pol III elongation complex have left large portions of the Rpc34 subunit in Pol III elongation complex concealed due largely to persisting challenges of spatially resolving a flexible part in macromolecular structures with cryo-EM^38–40^, particularly when the part is highly dynamic or associated with short-lived structural intermediate, as is the case of Rpc34 tWH domain during Pol III elongation. By engineering an azide-bearing UAA to Rpc34 WH2 and overcoming challenging issues associated with cross-activity with Bertozzi click chemistry, we achieved selective dye labelling of the UAA and measures smFRET between the acceptor on Rpc34 WH2 and the donor at a site in DNA. Surprisingly, the smFRET results show discrete multi-modal distribution of FRET efficiency, instead of continuous spectrum as anticipated for a highly dynamic part. By using nano-positioning analysis based on the smFRET distance constraints, we determined two credible volumes flanking Rpc34 WH2 to localize the multiple docking sites of Rpc34 WH2 on Pol III EC (Fig. 5a). Remarkably, in nearly 50% of Pol III EC molecules, Rpc34 WH2 could stochastically switch docking sites in real-time. This change of configuration may not require massive scale re-arrangement of Rpc34 subunit as it can be well accommodated by conformational change of the flexible linker connecting Rpc34 WH2 and WH3 domain, with the latter firmly anchoring on Pol III. Further dwell-time analysis of the state-switching Pol III EC reveals that Rpc34 WH2 exhibits site-specific characteristic life-times, where the short time-scale indicates the docking interactions are weak and promiscuous. It is well noticed that the docking life-times are anti-correlated with the distances between Rpc34 WH2 and WH3 (Fig. 5a).

**Fig. 5.**
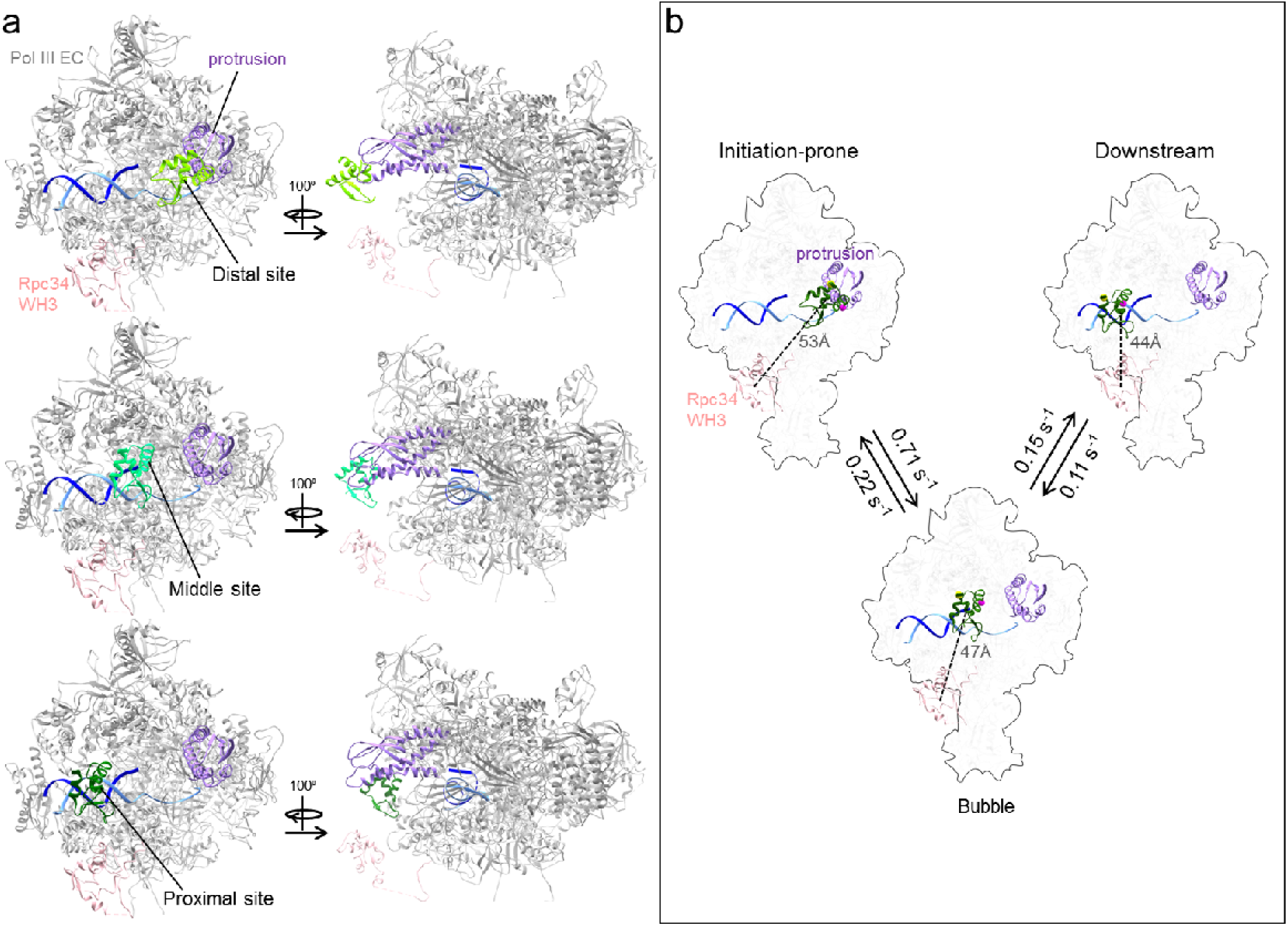
**Rpc34 WH2 Docking Sites in Pol III EC and the Pol III Configuration-Sampling Model.** (**a**) The three docking sites (distal, middle and proximal sites) of Rpc34 WH2 are localized according to the NPS credible volumes of L92 and K126 that flank Rpc34 WH2 (Fig. 4d). (**b**) In the Pol III configuration-sampling model, the spatial locations of the docking sites are the same as in (**a**), and the docking kinetics are derived from the dwell-time analysis and transition density plots (Fig. 3b & c). ***Upper left***: The initiation-prone (distal) site, based on the low-FRET state, is localized at the entrance of DNA-binding cleft, adjacent to the Rpc2 (Rpc 128) protrusion domain (light purple), resembling the position of Rpc34 WH2 (dark green) in the Pol III initiation complex. ***Lower***: The bubble (middle) site, derived from the middle-FRET state, is situated near the downstream end of the DNA bubble. ***Upper right***: The downstream (proximal) site, corresponding to the high-FRET state, is located in a region of downstream DNA. The transition rates among these states are obtained from the dwell-time analysis of state-change traces (Fig. 3 & Supplementary Fig. 8). The linker connecting Rpc34 WH2 and WH3 comprises 16 amino acids and can well accommodate the distances of 44 Å, 47 Å, and 53 Å observed between these two domains. Notably, the dwell-times of Rpc34 WH2 in Pol III EC are anti- correlated with the distances between Rpc34 WH2 and WH3.

We provisionally termed these docking sites as proximal, middle, and distal sites in reference to the downstream DNA or Rpc34 WH3 (Fig. 5a). For those active state-switching Pol III ECs, we propose a Pol III configuration-sampling model during elongation (Fig. 5b)—

in this model, Rpc34 tWH in a transcribing Pol III may frequently sample different docking sites so as to engage different functions. While dwelling on the proximal or middle site, not reported in previous structural studies, Rpc34 WH2 is situated above the DNA-binding channel that it may directly interact with the DNA^41,42^, perhaps contributing to holding the downstream DNA template in the DNA-binding cleft^16,43^, and to keeping the bubble in a melted state, respectively, for enhancing Pol III EC stability or processivity. These functional assignments are in line with the notion that Rpc34 tWH serves as a central protein module to integrate a dynamic protein-nucleic acid network during elongation established through function analysis^11^. What could be then the potential function of Rpc34 WH2 at the distal site?—a site that has a good overlap with that being occupied during initiation. Occupying a site of initiation during the stage of elongation seems to be counter-intuitive. However, in light of frequent re-initiation being a property unique to Pol III, by which a terminating enzyme can readily restart transcription on the same gene, the finding of the distal site may suggest a hypothetical but novel priming mechanism of re-initiation, orchestrated by Pol III adopting a configuration readily to engage with an initiation factor TFIIIB so as to reduce the kinetic barrier for forming the initiation complex at a promoter. With provisional assignment of the function at each site, we referred to distal, middle and proximal sites, as “initiation- prone”, “bubble” and “downstream” sites. Taken together, the findings of Rpc34 WH2 dynamic positioning allows one to postulate a function per docking position, conferring a structural basis for rationalizing Rpc34 tWH’s multi-functionality.

Winged-helix (WH) domain is a frequent recurring motif in TFIIE- or TFIIF-related sub-complexes within Pol I or Pol III, and its presence is to primarily mediate protein-protein interactions in intricate protein networks^44,45^. Our study provides plausible evidence that Rpc34 tandem winged-helix (tWH) domain in Pol III can interact with nucleic acids^41,42^ in addition to proteins, albeit in a weak and promiscuous manner—traits typically associated with intrinsically disordered proteins, which facilitate interactions with multiple partners to serve as molecular hubs within protein interaction networks^46^. In Pol I, the tWH domain is hosted by the Rpa49 subunit within the TFIIF-related sub-complex, which remains stably associated with Pol I. Rpa49 tWH seems to be flexible as its density could not be revealed by X-ray crystallography, but was identified in cryo-EM structures of various transcription stages^47^. These cryo-EM snapshot structures demonstrate that this tWH domain can shift to different positions to fulfil distinct functions. In an open complex (OC2), Rpa49 tWH binds to upstream DNA for promoter recognition or opening, and as transcription progresses, it shifts to a new position to displace Rrn3, facilitating Pol I’s escape from the promoter. The structural findings for Rpa49 tWH domain in Pol I^47^ and those for Pol III Rpc34 WH2 in the present study, together illustrate tWH’s capacity in mediating promiscuous interactions, a crucial feature underpinning the multi-functionality. Interestingly, cytosolic Pol III was discovered to be involved in immune surveillance against infection of varicella zoster virus (VZV) with severity associated with patient mutations in the tWH^48^ of hRPC6/hRPC39 (the human homologue of yeast Rpc34). Assuming that hRRC6/RPC39 tWH is likewise a tethering structural module that can dynamically and promiscuously interacting with protein-nucleic acid network, it is tempting to speculate that tWH in human Pol III would the part readily reach out to engage virus DNA^48^, which warrants a further study.

Finally, in this work we have implemented a simple thiol-capping scheme to tackle previously overlooked cross-reactivity issue associated with the strain-promoted azide-alkyne click reaction. The cross-reactivity of alkyne to thiol moiety has hindered this click chemistry from selectively labelling the azido moiety engineered into a protein unless this protein is without a cysteine or has no reactive cysteine. Prior to our work, Staudinger ligation has been opted to achieve selective labelling of an azido moiety engineered into a protein^49,50^. Thus, our scheme that can ensure selective strain-promoted azide-alkyne click reaction provides a useful tool for rendering large or native protein assemblies more amenable to smFRET-based dynamical structural biology^51^, pertinent to exploring protein dynamics underpinning the functions. Using this approach, we uncover dynamical positioning for a component in a stably and permanently associated sub-complex within a large protein complex. In the case of Rpc34 tWH, its positioning can vary in a function-dependent manner, which is previously unclear. This notion of a stably associated component being mobile might be further augmented by recent analyses of perplexing crosslinking data from holo-and transcribing Pol III^52^ that revealed Rpc82, the largest subunit in the TFIIE-related sub-complex, can undergo significant rearrangements within Pol III.

In conclusion, this study unravels a dynamic architecture of Pol III elongation apparatus in real-time for the first time, providing previously inaccessible spatial and temporal information for rationalizing the multi-functionality of a hub domain in Pol III transcription, furthering our understanding of Pol III machinery with mechanistic insights into Pol III transcription dynamics.

## Methods

### Plasmids, yeast strains and cell growth

Incorporation of UAA into yeast was achieved as described previously^19,20^. In brief, to generate a yeast strain in which UAA had been incorporated into Rpc34, the coding gene sequence of Rpc34 with a V5 epitope tag at the N terminus was cloned into the 2-micron vector pRS425 (LEU2+) (Fig. 1a). The TAG (amber) nonsense codon was introduced by in vitro mutagenesis into Rpc34 of the Rpc34/pRS425 plasmid to express a series of Rpc34 mutants in which UAA was incorporated at specific single positions. To incorporate an azido-UAA (4-Azido-L-phenylalanine, AzF), the plasmid pLH157 (tRNA_CUA_/BPA-tRNA synthetase) was modified to become pLH157-AzF (tRNA_CUA_/AzF- tRNA synthetase plasmid), in which the EcTyrRS gene was mutated to host amino acid changes Tyr37Leu, Asp182Ser, Phe183Met, and Leu186Ala according to the original gene mutations as previously described^19^. The Rpc34 mutant/pRS425 plasmid and pLH157-AzF were co-transfected into the *Saccharomyces cerevisiae* YLy3 yeast strain [MATα ade2::his3G his3Δ200 leu2Δ met15Δ lys2Δ trp1Δ63 ura3Δ (rpc34::KanMX4) Rpc34/pRS316 (Ura3+)] modified from the BY4705 strain. In this modified yeast strain, the chromosomal Rpc34 gene is disrupted by a KanMx gene cassette resistant to the drug G418. Additionally, the C terminus of the Rpc2 (Rpc128) protein hosts a TAP tag (FLAG3-His6-TEV cleavage sequence-protein A).

### Protein purification

The Pol III was purified as described previously^11^. In brief, yeast cell cultures were grown in YPD medium with 0.2 mM AzF to OD = 2 and then harvested. Cells from 6-liter cell cultures were pooled and re-suspended in the TAP-tag purification buffer containing 40 mM HEPES (pH 7.5), 350 mM NaCl, 10% glycerol, 0.1% Tween-20, 0.5 mM ethylenediaminetetraacetic acid (EDTA) and 1x protease inhibitors (1 μM pepstatin A, 1 mM phenylmethylsulfonyl fluoride (PMSF), 2.58 mM benzamidine and 0.7 mM leupeptin) and then lysed by glass bead beating. The cell lysate was centrifuged at 5,000 rpm with a JLA-8.1 rotor (Beckman Coulter) for 10 min, and ultra-centrifuged at 35,000 rpm with a Ti-45 rotor (Beckman Coulter) for 1 hour to collect the clarified lysate. The resulting supernatant was incubated overnight with 2 ml IgG-Sepharose resins (GE Healthcare) at 4°C. Subsequently, the bound proteins on the resins were washed once with 50 ml TAP-tag purification buffer and then re-suspended in 1 mL TEV cleavage buffer containing 10 mM Tris (pH 8.0), 150 mM NaCl, 10% glycerol, 0.1% NP-40, 0.5 mM EDTA, supplemented with 50 μg TEV protease and digestion was conducted overnight at 4°C. The eluted protein was verified by Coomassie-blue stained SDS-PAGE and Western blotting for the FLAG and V5 tags, aliquoted and stored at – 80 °C.

For *Saccharomyces cerevisiae* Maf1, the gene was codon optimized and synthesized in a pUC57 vector (GenScript Biotech.), and then sub-cloned into pDuet2 vector with the BamHI and XhoI restriction sites to produce a pDuet2-Maf1 plasmid. In this plasmid, the N-terminus of Maf1 has a

His6-HA-SUMO tag and the C-terminus has a TAP tag (FLAG-Twin Strep Tag with a HRV3C cleavage site in between to facilitate purification. To over-express this recombinant *S. cerevisiae* Maf1 in E.coli, this plasmid was transformed into Rosetta 2(DE3) competent cells (Merck/YB Biotech). Cells were first cultured in 2-liter LB medium containing 30 μg/ml chloramphenicol and 100 μg/ml Ampicillin at 37°C till OD_600_ reached 0.4 - 0.5, and the protein over-expression was induced with 0.1 mM IPTG overnight at 18°C. The cells were harvested by centrifugation with a JLA

8.1 rotor with 6,545 g for 30 minutes, and re-suspended in lysis buffer containing 1x PBS (pH 8.0), 1x protease inhibitor (made from 50x Coctail protease inhibitor, Roche), 5mM beta-mercaptoethanol, and then lysed by the constant cell disruption system (TS 2.2Kw, Constant Cell Disruption System). The whole cell extract was ultra-centrifuged at 36,000 rpm to achieve 188,000 g with a Ti-45 rotor for 30 minutes at 4°C to remove cell debris. The collected supernatant of 50 ml was first incubated with SUMO protease (ThermoFisher) overnight at 4°C to digest the SUMO tag, then mixed with 0.2 ml 50

% slurry of Strep-Tactin resin (ThermoFisher) and incubated at 4°C for 2 hours. After batch washing the Strep-Tactin resins by 100 column volume of “strep-wash” buffer (25 mM Tris-HCl pH 8.0, 500 mM NaCl, 1% NP-40 and 5 mM beta-mercaptoethanol) and equilibrating with an elution buffer (25mM Tris-HCl pH 8.0, 150 mM NaCl, 5% glycerol), the resins was incubated with the elution buffer supplemented with 100 μl of HRV3C with 1mM DTT for overnight at 4°C. Maf1 protein was eluted as fractions with 0.05 ml step; the fractions were further incubated with NTA resins for cleaning the HRV3C protease, examined by SDS-PAGE gel with Coomassie-Blue stain, and verified by MALDI-MS analysis. Two fractions with highest concentrations were pooled and exchanged to a storage buffer the same as elution buffer except 1mM DDT replaced by 10 mM TCEP, aliquoted and stored at – 80 °C.

### Pol III DIBO-labelling and EC formation

Purified Pol III was incubated with a DNA/RNA oligonucleotide scaffold—composed of TAMRA dye-labeled template DNA (T-DNA), biotinylated non-template DNA (NT-DNA) and 10-nucleotide (nt) RNA—to form the Pol III EC in buffer (100 mM HEPES pH 7.9, 400 mM KCl, 10% glycerol, 25 mM MgCl_2_, 5 mM EDTA). The oligonucleotide sequences and modifications are: (i) template strand with TAMRA dye labelling at the +7 position (5’-CATAAAAAACCCAAAAAAAGAGAGTATT-TAMRA dye-AATTGTTGAAGAAAGAGTATACTACATA); (ii) non-template strand (5’-biotin-TATGTAGATATGAGAAAGAAGTACAATTAAATACTCTCTTTTTTTGGGTTTTTT

ATG); (iii) RNA strand (5’-UCUUUCUUCA-3’). The three oligonucleotide strands were mixed and annealed to form the DNA/RNA scaffold with a 15-nucleotide bubble. After Pol III EC formation, free sulfhydryls of cysteines in the Pol III EC were capped with 0.2 mM N-ethylmaleimide (NEM) or 20 μM methyl-methanethiosulfonate (MMTS) as those concentrations were found to not to interfere with formation of Pol III EC, where high concentrations of NEM or MMTS did interfere with

formation of the Pol III EC as evidenced by disappearance of the respective band (>720 kD) in the native gel (Supplementary Fig. 2). Subsequently, 10 μM Alexa647-DIBO (C10408, Thermo-Fisher) was added into 3 μg of Pol III EC to specifically label AzF by copper-free click chemistry reaction at

4 °C overnight. Following the reaction, a desalting spin column (Roche) was used to remove excessive dye-DIBO reagent. Dye labelling was verified by fluorescence imaging of the SDS-PAGE gel, and EC formation was verified by fluorescence and Coomassie-blue-stained imaging of native PAGE (Thermo Fisher) (Supplementary Fig. 2). An elongation assay of Pol III EC was performed to ensure DIBO-dye labelling or cysteines protected by NEM or MMTS did not affect Pol III elongation activity (data not shown).

### SmFRET system equipped with alternating laser excitation

Our smFRET experiments and analytical methodology were conducted as described previously^53^. In brief, approximately 100 pM Pol III EC labeled with the donor-acceptor dye pair was immobilized on a clean cover-glass that was surface-coated with Neutravidin (Thermo Fisher) and biotin-PEG (Laysan Bio) in a sample chamber^54^. The oxygen scavenger imaging buffer (175 nM protocatechuate- 3,4-dioxygenase, 7.8 mM protocatechuic acid, 2 mM Trolox, 50 mM Tris:HCl pH7.8, 150 mM NaCl, 1 mM PMSF and 0.1 mg/ml BSA) was injected into the chamber to increase fluorophore photo- stability^55,56^. Wide-field fluorescence images were acquired using a custom-built dual-view total internal reflection fluorescence (TIRF) microscope in which a 532 nm-wavelength laser (DPGL-2100, Photop Technologies) and a 638 nm-wavelength laser (LuxX 638–100, Omicron) were used to excite the donor and acceptor dyes; an oil-immersion objective (UPLSAPO100XO, Olympus, NA 1.49) was used to focus the laser beam onto the sample and collect fluorescence. The dichromatic mirror (FF640-FDi01, Semrock) separated the collected fluorescence photons into the donor and the acceptor emission arms, with the two different color images projected onto different areas of the EMCCD camera (DV887DCS-BV, Andor Technology). The movies were captured with a frame rate of 5fps, namely 200 milliseconds per frame. In addition, we implemented an alternating laser excitation (ALEX)^57^ scheme using a shutter system (LS3ZM2, Uniblitz Electronic) and relay circuit for switching between the excitation of the donor and that of the acceptor as described^58^. ALEX imaging filters out the time periods when the donor or acceptor is absent due to switching to unwanted photo- physical states.

### Simulation of dye volume and the predicted FRET efficiency

In order to predict the FRET efficiencies anticipated from the positions of Rpc34 WH2 in different existing structural models, we computationally placed the acceptor at the targeted amino acid in Rpc34 WH2 based on a structural model and the donor dye at a site of DNA modeled into it. The modeling of DNA into a structural model was performed by aligning a particular structural model

against Pol III EC (PDB: 5fj8) based on Rpc 160 (Rpc1) (Supplementary Fig. 1). Accessible volumes of dyes were simulated by FPS software^31^ based on donor dye at DNA and acceptor dye at Rpc34 WH2 with designated dye parameters (size of dye and the length and width of linker) (Supplementary Table 2). The distances R_MP_ between the mean dye positions of two accessible volumes of FRET pair in different models were obtained. We calculated the predicted FRET efficiencies for different models based on the mean dye distances and the Förster radius of dye pair (Supplementary Table 3 & 4). In addition, in the simulation of dye accessible volume attached to a benchmark DNA^35^ where the two nucleotides labeled by dye were separated by 15 bp. (Supplementary Fig. 3), the double-strand DNA model was built up using the web server of Web 3DNA^59^. The smFRET measurement of double-labelled benchmark DNA^35^ demonstrates the precision of our system (Supplementary Fig. 3).

### SmFRET data analysis and dwell-time analysis and exponential fitting

We used a custom-built two-channel TIRF system to record fluorescence images of the donor and acceptor^53^. Fluorescence movie data were analysed for identifying the co-localized donor/acceptor pair and extracting the donor and acceptor intensities as time trajectories using algorithms and programs as described^60,61^. The extracted signal intensities of the donor and acceptor were used to calculate FRET efficiencies according to the following equation:

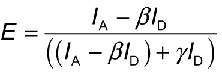

as described^34^ and according to the method of individual γ normalization^34^. *I*_A_ and *I*_D_ represent the measured intensities of the acceptor and donor, respectively; β*I*_D_ corrects for leakage of donor emission into the acceptor channel, and γ corrects for the difference in detection efficiency and quantum yield between the donor and acceptor. We adopted acceptor-photo-bleaching FRET^34^ to validate each candidate FRET trace and also to obtain the absolute FRET efficiencies. This method assesses a time trace based on anti-correlated donor/acceptor signals at the time-point when the acceptor is photo-bleached. Using the stringent criterion of acceptor-photo-bleaching^34^, we could screen candidate smFRET traces selected based on donor/acceptor co-localization. Typically, we obtained one to two authentic smFRET traces from each movie. Moreover, considering that the acceptor may not always be active in the time trace of a single molecule, we excited both channels using ALEX to continuously survey the presence of an acceptor so as to detect the period when it is not in the active state, namely in the dark state, prior to photo-bleaching. Using this approach, we filtered out those artefact traces due to acceptor quenching not due to FRET. We used the frame-wise FRET efficiency to build a histogram and fitted it using multiple Gaussian distributions for extracting the mean FRET efficiencies. Alternatively, the mean FRET efficiencies were obtained from stationary traces by smoothing a trace using a denoising algorithm based on wavelets and Bayesian inference^36^, and the mean FRET efficiencies were used to categorize a stationary trace. A frame-wise FRET efficiency histogram was built from time traces in a category, and was fitted with single Gaussian

distribution for extracting the mean efficiency. These two approaches gave identical results in terms of the number of states and the centers of Gaussian distributions. The levels and periods of different states in inter-state converting traces were identified by hidden Markov modeling (HMM)^37^, and the frame data in the same level was pooled to build a histogram and fitted with one or two exponential function(s) in the dwell-time analysis. The transitions between states in the FRET time traces were used to build the transition density plot (Fig.3c).

### Nano-positioning system analysis

Nano-positioning system (NPS) analysis was performed by using the Fast-NPS method^62^. The credible volume accessible by the antenna dye (acceptor) was calculated using the Fast-NPS software, with the parameters of the mean and standard deviation of FRET efficiencies derived from the smFRET histogram, the coordinates of the satellite dye (donor) attached to DNA, the Förster radius of the dye pair, fluorescence anisotropy of both dyes, and the properties of the dye and linker (dye diameter: 13 Å; linker length: 13 Å; linker diameter: 4.5 Å, as suggested in Ref. 62). The coordinates of the satellite dye attachment point were based on the atomic model of Pol III EC (PDB: 5fj8). The Förster radius of the dye pair is 6.0 nm^32,33^. The anisotropy of the donor dye on DNA was measured as

∼0.3 using a lifetime technique^35^ by means of a time-resolved laser scanning confocal microscope (Q2, ISS, USA) with the measured value in good agreement with those of previous studies^35^. For the anisotropy of the acceptor dye on UAA in Pol III, we adopted the value of ∼0.3 reported by the Deniz group^24^.

## Data availability

The raw FRET movies or time traces data in this study are available from the corresponding authors upon reasonable request.

## Acknowledgements

W.H.C. is indebted to Dr. Jiun-Jie Shie of Institute of Chemistry, Academia Sinica for commenting on the limitations of Bertozzi reactions for bio-orthogonal applications. The authors thank John O’Brien for initial English editing. This work is supported by Academia Sinica [AS-TP-110-L11], and Ministry of Science and Technology (MOST) of Taiwan [105-2627-M-001-010, 106-2627-M-001- 009 and 107-2627-M-001-005] to W.H.C. and H.T.C. J.S.W. has been supported by an Academia Sinica Postdoctoral Fellowship. I.P.T. is supported by an Academia Sinica Investigator Award [AS- IA-110-M05].

## Author contributions

W.H.C. and H.T.C. conceived the project. J.S. W. performed dye attachment simulation, single-molecule experiments and analysis. Y.C.L. (Yu-Chun Lin) and Y.Y.W. performed the construction of UAA yeast strains and yeast Pol III purification with bulk biochemistry. J.S.W. and Y.Y.W. screened

thiol-blocking reagents and optimized conditions to achieve selective Bertozzi azido-UAA labelling.

H.H.L. and I.P.T. developed efficient algorithms for streamlining the single-molecule analysis. Y.C.L. (Yang-Chih Liu) conducted the purification of Maf1 used for the control experiments. W.H.C. and

J.W.C. designed the UAA labelling scheme. J.S.W., H.T.C., and W.H.C. wrote the manuscript, with

I.P.T. contributing to further editing.

## Additional Information

### Supplementary Information

Supplementary information is available in a separate file

### Correspondence and requests for materials

should be addressed to Wei-Hau Chang.

