## Supplementary Information for "Structural underpinning of multi-functionality of TFIIE-related tandem-winged-helix in RNA polymerase III"

### for

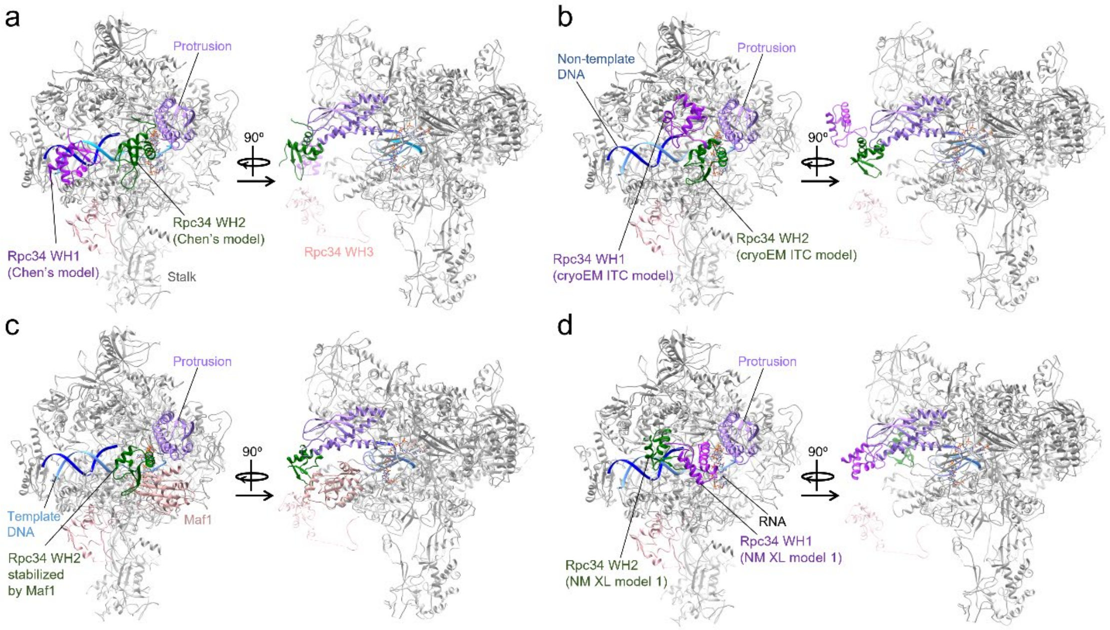


e

| Structural model | Distances between the mean dye positions $R_{MP}$ (Donor: T-DNA(+7), Acceptor: Rpc34-K126) (Å) | Predicted FRET efficiency |
|---|---|---|
| PA-XL | 72.5 | 0.21 |
| Cryo-EM ITC | 70.4 | 0.24 |
| Pol III with Maf1 | 68.9 | 0.26 |
| NM XL model 1 | 46.2 | 0.80 |

**Supplementary Fig. 1 │ Positions of Rpc34 tandem-winged-helix (tWH) domain in various models of Pol III in different contexts from literature.** (**a**) Rpc34 WH2 (dark green) in Chen's photo-activated crosslinking (PA-XL) model[12], (**b**) Rpc34 WH2 in the initial transcribing complex (ITC, PDB: 6f41)[13], (**c**) Rpc34 WH2 stabilized by Maf1 (PDB: 6tut)[15], and (**d**) Rpc34 WH2 in Nilges & Muller crosslinking (NM-XL) Model 1[52]. All models were docked into Pol III elongation complex model (gray, PDB: 5fj8) based on alignment of Rpc1 between models. (**e**) The distances between the mean dye positions $R_{MP}$ and the predicted FRET efficiency in various Pol III models. The donor dye and acceptor are labeled at T-DNA(+7) and Rpc34-K126, respectively.

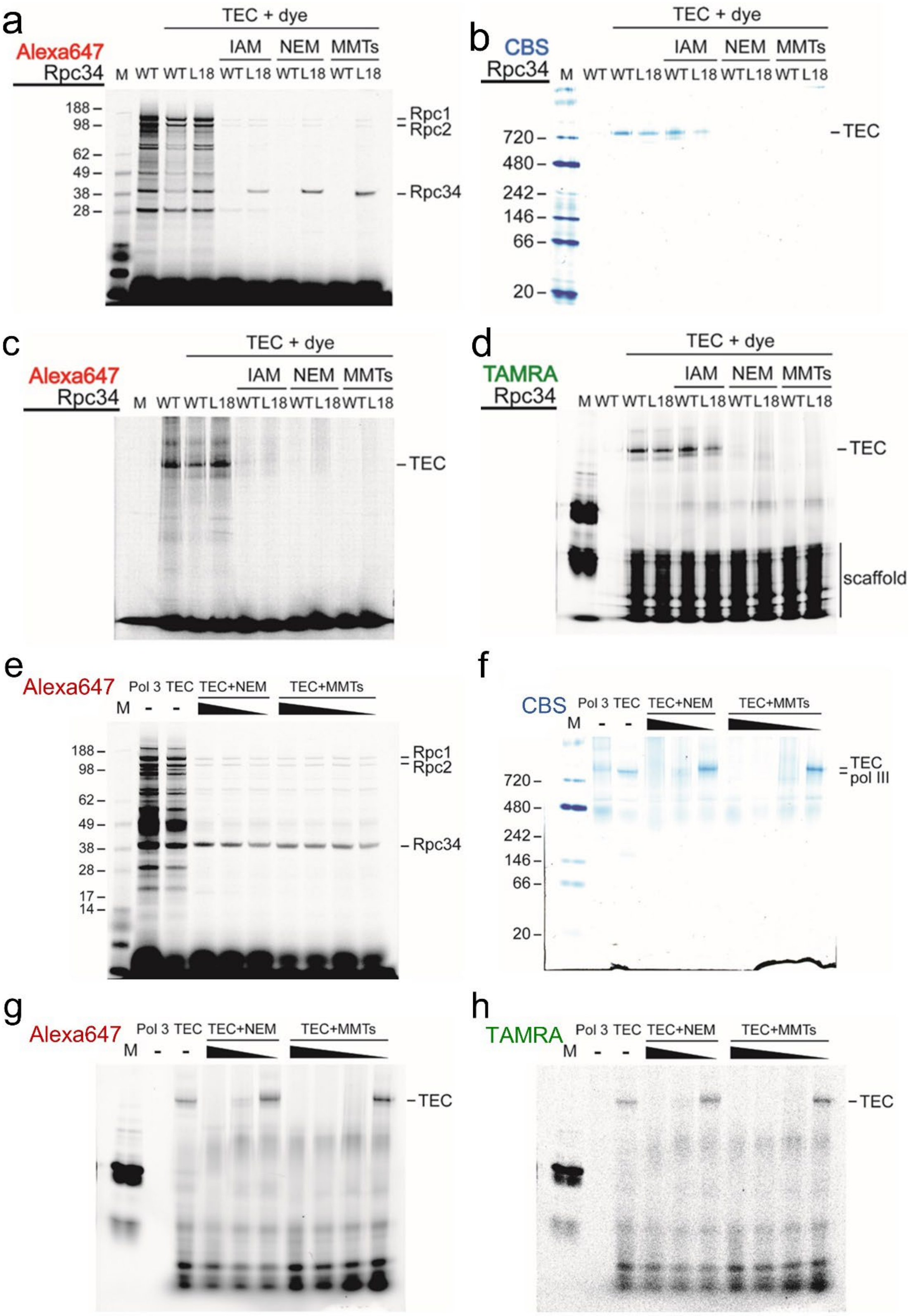

a
Alexa647
Rpc34
TEC + dye
IAM NEM MMTs
M WT WT L18 WT L18 WT L18 WT L18
188
98
62
49
38
28
Rpc1
Rpc2
Rpc34
b
CBS
Rpc34
TEC + dye
IAM NEM MMTs
M WT WT L18 WT L18 WT L18 WT L18
720
480
242
146
66
20
TEC
c
Alexa647
Rpc34
TEC + dye
IAM NEM MMTs
M WT WT L18 WT L18 WT L18 WT L18
TEC
d
TAMRA
Rpc34
TEC + dye
IAM NEM MMTs
M WT WT L18 WT L18 WT L18 WT L18
TEC
scaffold
e
Alexa647
M Pol 3 TEC TEC+NEM TEC+MMTs
- -
188
98
62
49
38
28
17
14
Rpc1
Rpc2
Rpc34
f
CBS
M Pol 3 TEC TEC+NEM TEC+MMTs
- -
720
480
242
146
66
20
TEC
pol III
g
Alexa647
M Pol 3 TEC TEC+NEM TEC+MMTs
- -
TEC
h
TAMRA
M Pol 3 TEC TEC+NEM TEC+MMTs
- -
TEC

**Supplementary Fig. 2 │ Selective labeling of incorporated azido-UAA by screening blockers that suppress cross-reactivity between cyclooctynes and thiols in cysteines.** (**a**)-(**d**) Thio-reactive molecules (iodoacetamide (IAM), N-ethylmaleimide (NEM) and S-methyl methanethiosulfonate (MMTS)) were tested to suppress non-specific dye labeling. The concentrations of three blockers are 20 mM. Pol III EC was labeled at the azido-UAA in Rpc34 L18 with Alexa647-DIBO and the template DNA was labeled with TAMRA. (**a**) Alexa647 fluorescence image of a SDS-PAGE gel of wild-type and UAA-substituted (Rpc34 L18) Pol III EC. Rpc1 and Rpc2 are the two largest subunits of Pol III and synonymous to Rpc160 and Rpc128, respectively. (**b**) Coomassie blue-stained (CBS) image, (**c**) Alexa647 image and (**d**) TAMRA image of a Native-PAGE gel of wild-type and UAA-substituted (Rpc34 L18) Pol III EC. (**e**)-(**h**) Two blockers were titrated to suppress non-specific dye labeling but permit Pol III EC formation. (**e**) Alexa-647 fluorescence image of a SDS-PAGE gel of Pol III EC with three NEM concentrations (20 mM, 2 mM and 0.2 mM) and four MMTS concentrations (20 mM, 2 mM, 0.2 mM and 20 μM). Note that NEM or MMTS at the lowest concentration (0.2 mM for NEM and 20 μM for MMTS) proved sufficient to reduce non-specific dye labeling. (**f**) CBS image, (**g**) Alexa647 image and (**h**) TAMRA image of a Native-PAGE gel of Pol III EC with three NEM concentrations (20 mM, 2 mM and 0.2 mM) and four MMTS concentrations (20 mM, 2 mM, 0.2 mM and 20 μM). Note that 0.2 mM NEM or 20 μM MMTS allowed formation of Pol III EC.

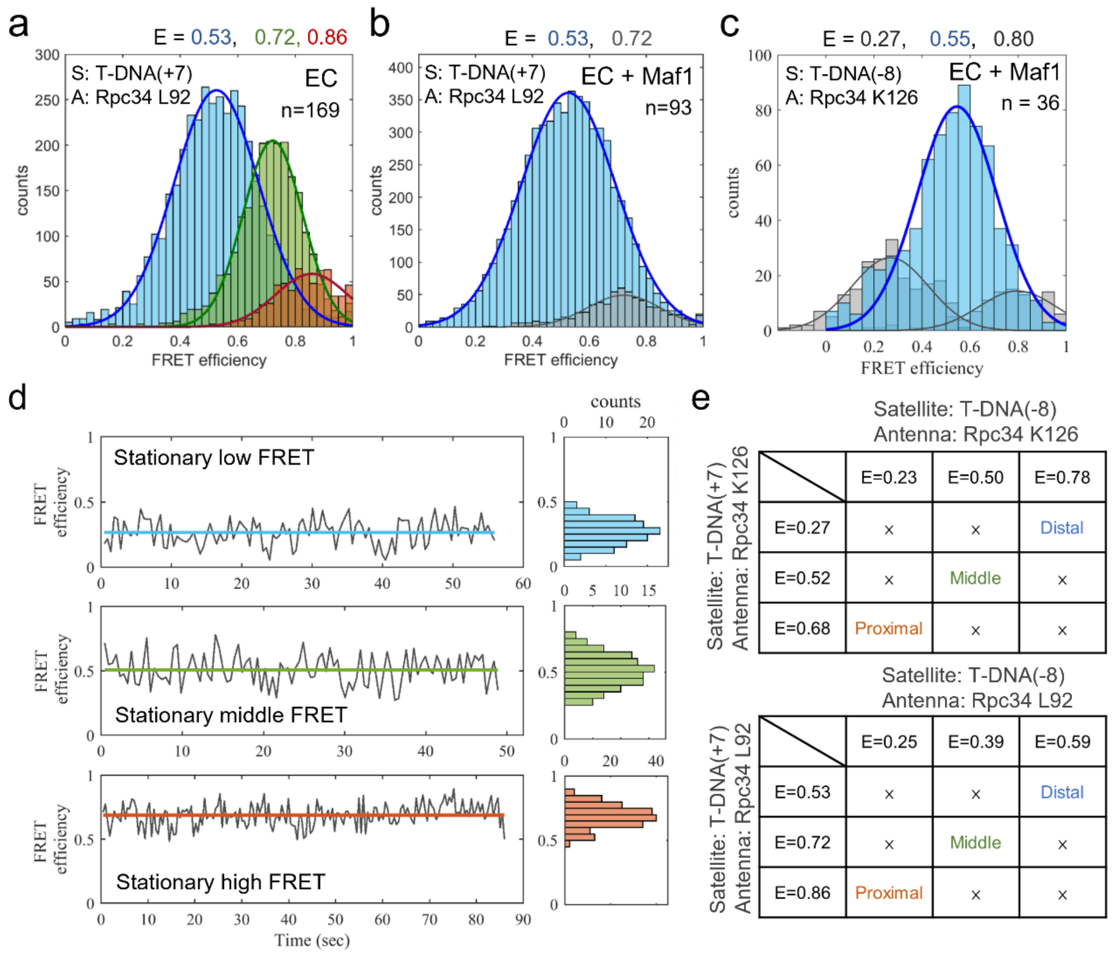


**Supplementary Fig. 3 │ FRET efficiency histograms of Rpc34 WH2 in Pol III EC without or with Maf1 and raw representative time trajectories of Rpc34 WH2 in Pol III EC.** (**a**) The FRET efficiency histogram of Rpc34 WH2 in Pol III EC probed by a donor-acceptor (D/A) pair at DNA(+7)/Rpc34 L92 (+7/92) (Same as the histogram in the bottom left in Fig. 4b). (**b**) The FRET efficiency histogram of Rpc34 WH2 in Pol III EC with Maf1 probed by D/A pair +7/92. The major population centered at E=0.53 indicates the FRET state of Maf1-stabilized Rpc34 WH2. E: the center of FRET efficiency distribution; D: donor; A: acceptor; n: number of molecules. (**b**) The FRET efficiency histogram of Rpc34 WH2 in Pol III EC with Maf1 probed by D/A pair -8/126. The FRET efficiency (E=0.55) of the major population matches well with the predicted FRET efficiency (E=0.54 as shown in Supplementary Table 3). (**c**) The raw representative time trajectories of stationary low-, middle- and high-FRET states of Rpc34 WH2 in Pol III EC (FRET pair +7/126). *Left*: The three lines (blue, green and orange) represent the mean FRET efficiencies. *Right*: The FRET efficiency histograms of corresponding to the traces of stationary low-, middle- and high-FRET states in the left panel. (**d**) The matrices of satellite-antenna pairs for NPS analysis where the antenna dye was labeled at K126 (upper panel) or L92 (bottom panel).

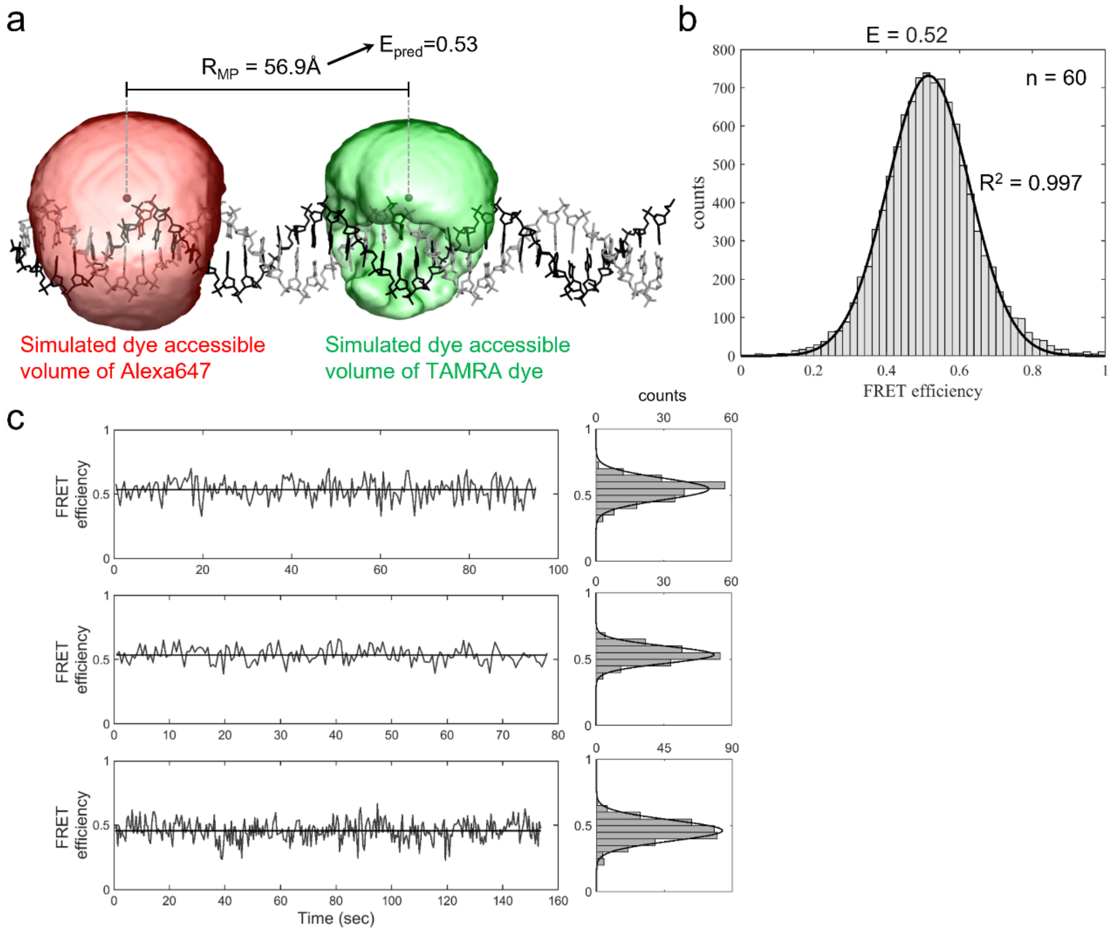


**Supplementary Fig. 4 │ Predicted FRET efficiency and smFRET measurement of double-labeled benchmark DNA.** (**a**) Dye accessible volumes of Alexa647 and TAMRA were simulated by FPS software[31]. The DNA sequence and the dye labeling positions of the benchmark DNA was the same as those in Ref. 35, where the two nucleotides labeled by dye were separated by 15 bp. The DNA model was built by using Web 3DNA[59]. The predicted FRET efficiency $E_{pred}$ was calculated from the distance between the centers of two dye accessible volumes $R_{MP}$. (**b**) The FRET efficiency histogram of double-labeled benchmark DNA fitted with single Gaussian function shows the center of distribution (E=0.52) agrees well with the predicted FRET efficiency (E=0.53). (**c**) *Left*: Three representative time trajectories of double-labeled benchmark DNA show stationary FRET level. *Right*: The FRET efficiency histograms of corresponding to the traces in the left panel. The distribution in each FRET efficiency histogram was best fitted with single Gaussian distribution.

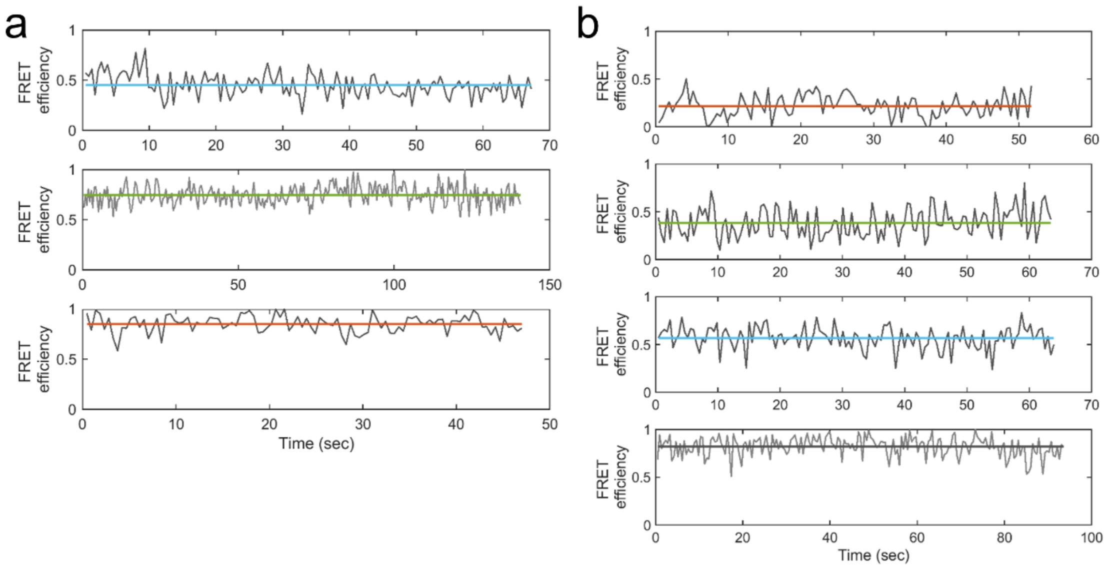


**Supplementary Fig. 5 │ The raw representative time trajectories of stationary FRET states of Rpc34 WH2 in Pol III EC, where the acceptor dye was labeled at Rpc34 L92.** (**a**) The three time trajectories correspond to the populations of E=0.53, E=0.72 and E=0.86 in the FRET efficiency histogram of Rpc34 WH2 in Pol III EC probed by donor-acceptor (D/A) pair +7/92 (bottom left in Fig. 4b). The three lines (blue, green and orange) represent the average FRET efficiencies of each time trajectory. (**b**) The four time trajectories correspond to the populations of E=0.25, E=0.39, E=0.59 and E=0.81 in the FRET efficiency histogram of Rpc34 WH2 in Pol III EC probed by D/A pair -8/92 (bottom right in Fig. 4b). The four lines (orange, green, blue and gray) represent the mean FRET efficiencies of each time trajectory.

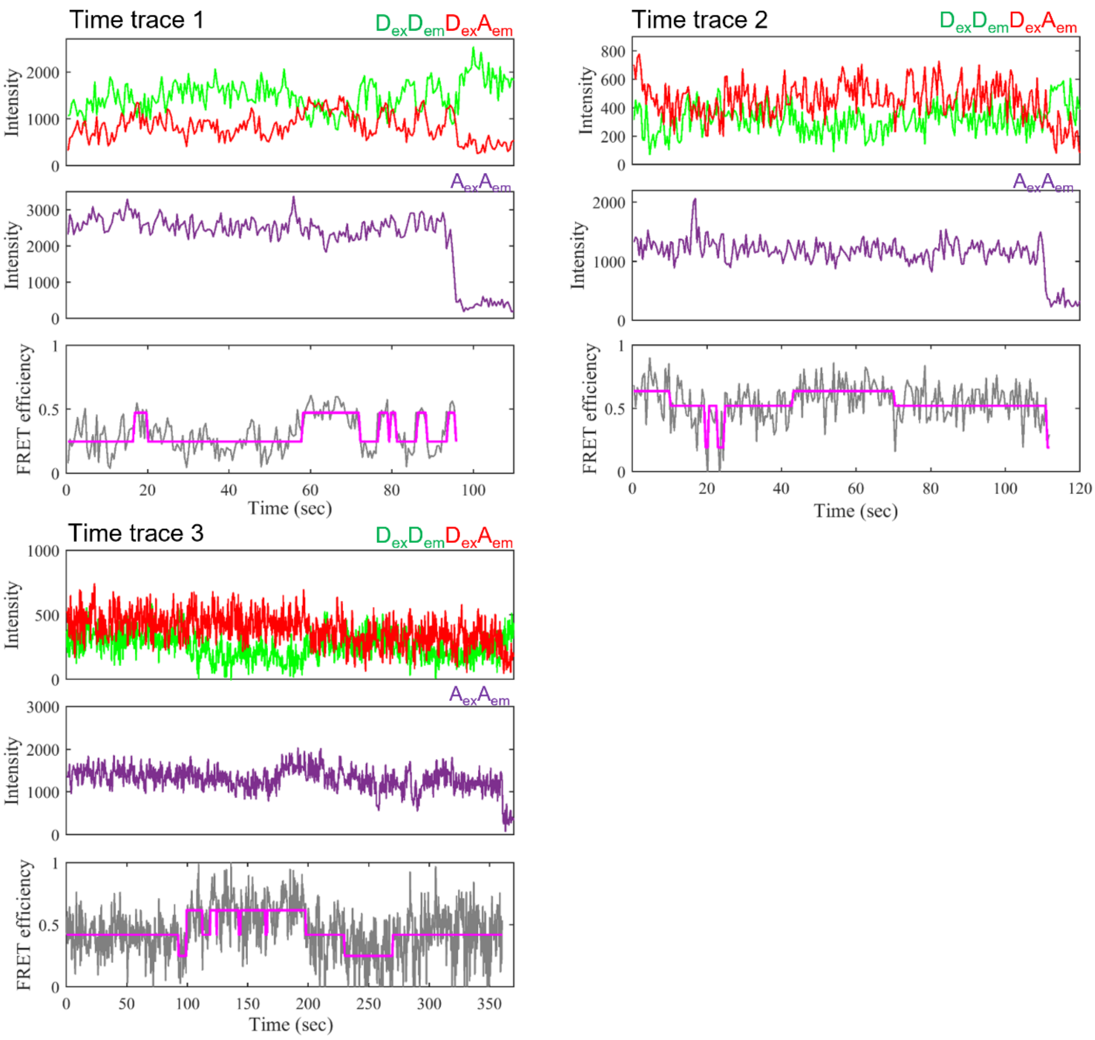


**Supplementary Fig. 6 │ Three representative time trajectories of inter-state switching FRET with ALEX channel (Time traces 1-3 in Fig. 3a)**. For each single-molecule time trace, three panels corresponding to three channels are shown. The upper panel represents the donor intensity ($D_{em}$), colored in green, when donor is excited using 532 nm laser light ($D_{ex}$), and the acceptor intensity ($A_{em}$), colored in red, also when donor is excited. The middle panel represents the acceptor intensity ($A_{em}$), colored in purple, when acceptor is excited using 638 nm laser light ($A_{ex}$). The lower panel represents the corresponding time traces of FRET efficiencies (gray trace) that are calculated from $D_{em}$ and $A_{em}$ from the upper panel. Hidden Markov modeling (HMM) is used for identifying the level of each FRET state, colored in magenta, and the respective durations.

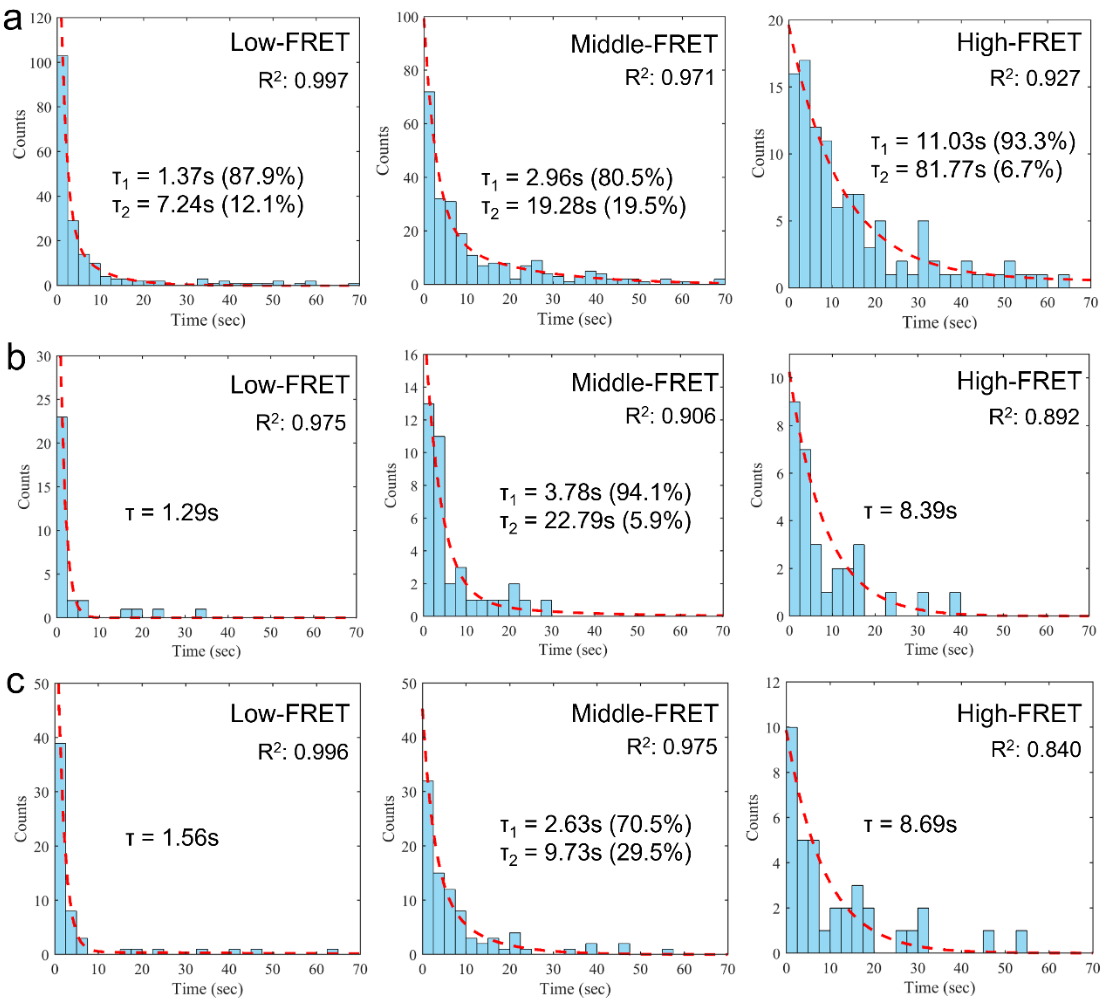


**Supplementary Fig. 7 │ The dwell time histograms for low-, middle-, and high-FRET efficiencies in the three individual smFRET measurements.** TAMRA was labeled at the +7 position of the template DNA. DIBO-Alexa647 was conjugated to Rpc34 K126 via Bertozzi strain-promoted azide-alkyne [3+2] cycloaddition (SpAAC). (a) The dwell time histogram was built from time durations extracted by HHM analysis from 62 inter-state switching time traces out of 110 time-traces in the first experiment. (b) The dwell time histogram was built from time durations extracted by HHM analysis from 25 inter-state switching time traces out of 80 time-traces in the second experiment. (c) The dwell time histogram was built from time durations extracted by HHM analysis from 18 inter-state switching time traces out of 54 time-traces in the third experiment. The dwell time histograms were well fitted with one or two exponential function(s). In the latter case, the exponential function with shorter life time (τ) always dominates. The average dwell times for low-, middle- and high-FRET are 1.41±0.11 sec, 3.12±0.48 sec and 9.37±1.18 sec, respectively.

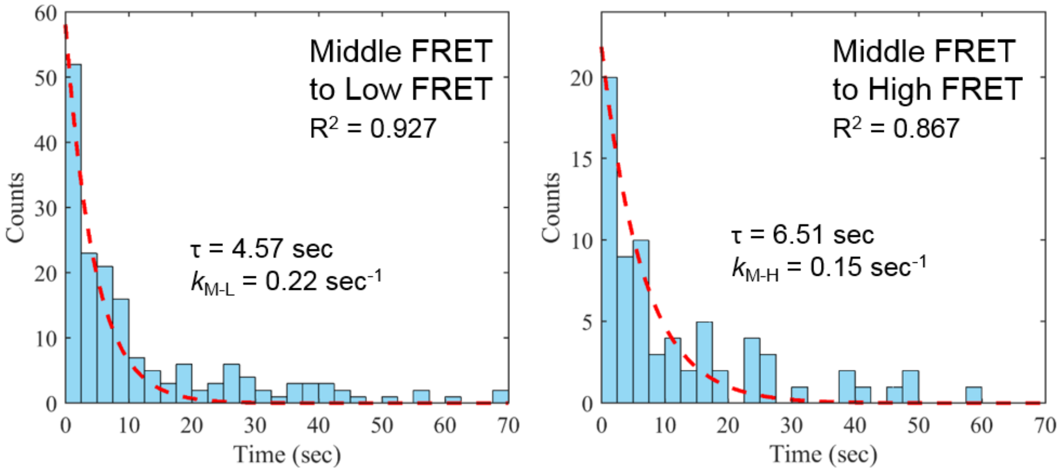


**Supplementary Fig. 8 │ The dwell time histograms for the middle to low FRET transition and the middle to high FRET transition.** The dwell time data for the middle FRET were separated into two groups based on the transition type: from the middle to low FRET, or from the middle to high FRET. Both histograms were best fitted to single exponential function. τ: dwell time; $k_{\text{M-L}}$: transition rate from middle to low FRET; $k_{\text{M-H}}$: transition rate from middle to high FRET.

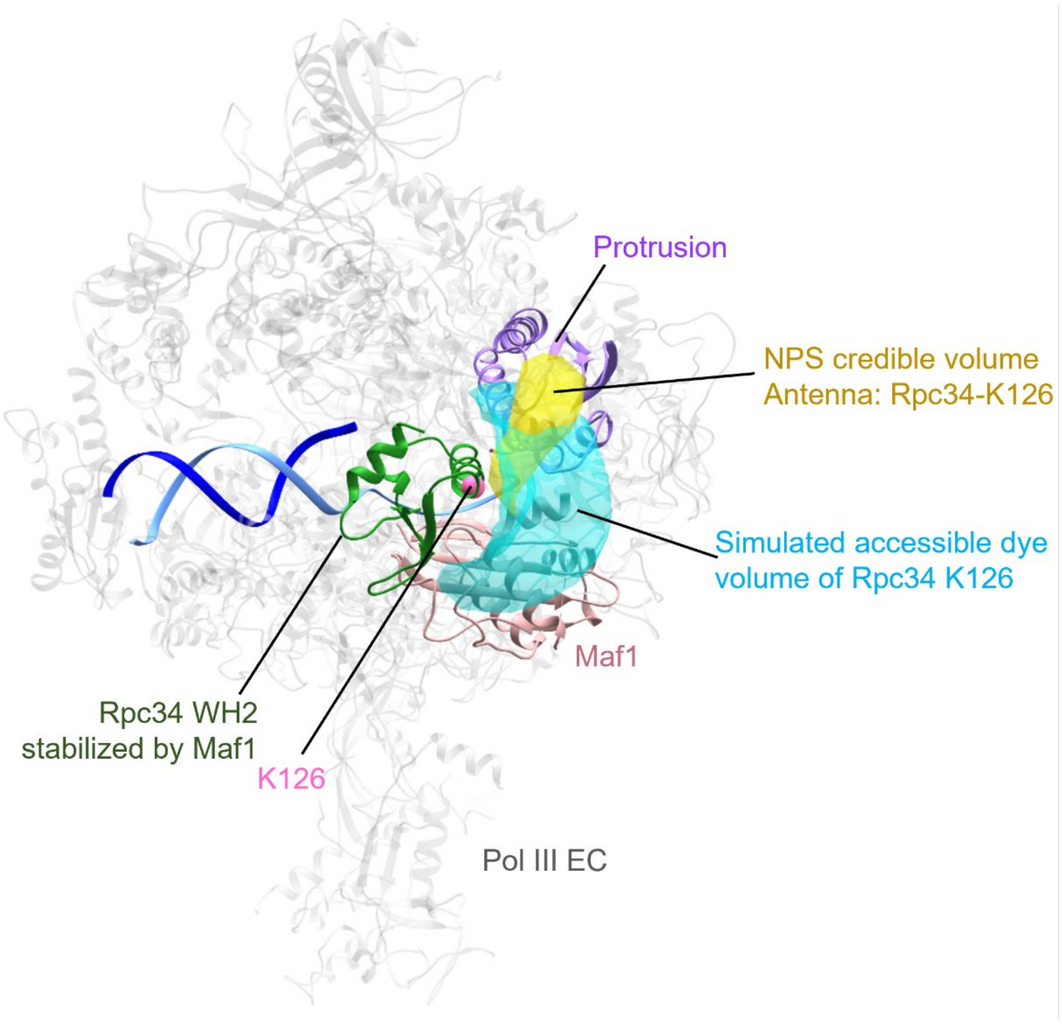


**Supplementary Fig. 9 │ NPS credible volume and simulated accessible dye volume of Maf1-stabilizaing Rpc34 WH2.** The antenna dye was labeled at Rpc34 K126 (hot pink sphere). The NPS credible volume (yellow) was analyzed by using two independent smFRET measurements of SA pair +7/126 and -8/126 in Pol III EC including Maf1 (Fig. 2c and Supplementary Fig. 3c). The accessible dye volume (cyan) of antenna dye was simulated by FPS software based on the Pol III EC model (PDB: 5fj8) docked with Rpc34 WH2 and Maf1 (PDB: 6tut) and dye parameters (size of dye and the length and width of linker) (Supplementary Table 2).

**Supplementary Table 1 │ Number of total and exposed cysteines on the Pol III surface**

| Subunit | Number of total cysteines | Number of exposed cysteines* (PDB: 5fj8) | Number of exposed cysteines within FRET sensing range |
|---|---|---|---|
| Rpc1(160) | 24 | 6 | 5 |
| Rpc2(128) | 19 | 4 | 3 |
| Rpc82 | 4 | 2 | 2 |
| Rpc53 | 0 | 0 | 0 |
| AC40 | 5 | 3 | 0 |
| Rpc37 | 0 | 0 | 0 |
| Rpc34 | 3 | 0 | 0 |
| Rpc31 | 0 | 0 | 0 |
| ABC27 | 3 | 2 | 2 |
| Rpc25 | 4 | 2 | 0 |
| Rpc17 | 1 | 0 | 0 |
| ABC14.5 | 2 | 1 | 0 |

*The exposed cysteines were determined by the property of relative amino acid exposure in UCSF Chimera (Ref: https://www.cgl.ucsf.edu/chimera/docs/UsersGuide/surfnorm.html) with a threshold of 0.6 (where the value of this property ranged from 0 to 1.2).

**Supplementary Table 2 │ The dye parameters for the simulation of dye accessible volumes.**

| Dye reagent for labeling | Length of linker, $L_{link}$ (Å) | Width of linker, $W_{link}$ (Å) | Dimension 1 of dye, $R_{dye(1)}$ (Å) | Dimension 2 of dye, $R_{dye(2)}$ (Å) | Dimension 3 of dye, $R_{dye(3)}$ (Å) |
|---|---|---|---|---|---|
| Alexa647-DIBO | 24.0 | 4.5 | 11.0 | 4.7 | 1.5 |
| TAMRA | 20.0 | 4.5 | 5.0 | 4.0 | 1.5 |
| DyLight650-phosphine | 22.5 | 4.5 | 11.0 | 4.7 | 1.5 |

*Dye parameters were suggested from the manual of FPS software and estimated from chemical structure of dye reagents.

**Supplementary Table 3 │ Predicted FRET efficiencies of FRET pair +7/126 and -8/126 in different structural models of Pol III.**

| Structural model | Distances between the mean dye positions $R_{MP}$ of pair +7/126* (Å) | Predicted FRET efficiency of pair +7/126† | Distances between the mean dye positions $R_{MP}$ of pair -8/126* (Å) | Predicted FRET efficiency of pair -8/126† |
|---|---|---|---|---|
| Pol III with Maf1[15] | 68.9 | 0.26 | 56.4 | 0.54 |
| Cryo-EM ITC[13] | 70.4 | 0.24 | 48.1 | 0.76 |
| Cryo-EM PIC (Muller)[13] | 69.6 | 0.25 | 47.9 | 0.76 |
| Cryo-EM PIC (Vannini)[14] | 67.1 | 0.30 | 45.5 | 0.81 |
| PA-XL[12] | 72.5 | 0.21 | 59.0 | 0.48 |
| NM XL model 1[44] | 46.2 | 0.80 | 57.9 | 0.50 |
| NM XL model 2[44] | 51.6 | 0.67 | 49.5 | 0.72 |
| NM XL model 3[44] | 46.5 | 0.79 | 43.7 | 0.85 |
| NM XL model 4[44] | 40.0 | 0.90 | 60.1 | 0.45 |

*The dye attached positions of donor and acceptor of FRET pair +7/126 are the atom C7 of +7 position in template DNA and the atom CD of Rpc34-K126, respectively. The dye attached positions of donor and acceptor of FRET pair -8/126 are the atom C5 of -8 position in template DNA and the atom CD of Rpc34-K126, respectively.

†The estimation of FRET efficiency is based on the Forster radius 58.1Å of the dye pair TAMRA and Alexa647.

**Supplementary Table 4 │ Predicted FRET efficiencies of FRET pair +7/92 and -8/92 in different structural models of Pol III.**

| Structural model | Distances between the mean dye positions $R_{MP}$ of pair +7/92* (Å) | Predicted FRET efficiency of pair +7/92† | Distances between the mean dye positions $R_{MP}$ of pair -8/92* (Å) | Predicted FRET efficiency of pair -8/92† |
|---|---|---|---|---|
| Pol III with Maf1[15] | 51.7 | 0.63 | 61.8 | 0.37 |
| Cryo-EM ITC[13] | 56.4 | 0.50 | 54.6 | 0.55 |
| Cryo-EM PIC (Muller)[13] | 59.9 | 0.42 | 57.9 | 0.47 |
| Cryo-EM PIC (Vannini)[14] | 60.9 | 0.39 | 60.4 | 0.40 |
| PA-XL[12] | 43.2 | 0.83 | 48.3 | 0.72 |
| NM XL model 1[44] | 32.2 | 0.97 | 71.2 | 0.20 |
| NM XL model 2[44] | 55.2 | 0.54 | 65.8 | 0.29 |
| NM XL model 3[44] | 64.4 | 0.32 | 69.6 | 0.22 |
| NM XL model 4[44] | 54.4 | 0.56 | 47.5 | 0.74 |

*The dye attached positions of donor and acceptor of FRET pair 3 are the atom C7 of +7 position in template DNA and the atom CG of Rpc34-L92, respectively. The dye attached positions of donor and acceptor of FRET pair 4 are the atom C5 of -8 position in template DNA and the atom CG of Rpc34-L92, respectively.

†The estimation of FRET efficiency is based on the Forster radius 56.6Å of the dye pair TAMRA and DyLight650.
